# Gene network modeling via TopNet reveals robust epistatic interactions between functionally diverse tumor critical mediator genes

**DOI:** 10.1101/2020.10.07.328542

**Authors:** Helene R. McMurray, Aslihan Ambeskovic, Laurel A. Newman, Jordan Aldersley, Vijaya Balakrishnan, Bradley Smith, Harry A. Stern, Hartmut Land, Matthew N. McCall

## Abstract

Malignant cell transformation and the underlying genomic scale reprogramming of gene expression require cooperation of multiple oncogenic mutations. Notably, this cooperation is reflected in the synergistic regulation of downstream genes, so-called cooperation response genes (CRGs). CRGs impact diverse hallmark features of cancer cells and are not known to be functionally connected. Yet, they act as critical mediators of the cancer phenotype at an unexpectedly high frequency of >50%, as indicated by genetic perturbations. Here we demonstrate that CRGs function within a network of strong genetic interdependencies that are critical to the robustness of the malignant state. Our approach, termed TopNet, utilizes attractor-based ternary network modeling that takes the novel approach of incorporating uncertainty in the underlying gene perturbation data and is capable of identifying non-linear gene interactions. TopNet reveals topological gene network architecture that effectively predicts previously unknown, functionally relevant epistatic gene interactions, and thus, among a broad range of applications, has utility for identification of non-mutant targets for cancer intervention.

## Introduction

Oncogenic mutations are critical for both cancer initiation and maintenance, which has driven efforts to inhibit targetable oncoproteins as an effective strategy for cancer treatment. Targetable mutations, however, are found only in a small fraction of cancers. Thus, broader strategies for cancer intervention need to be developed, such as targeting non-mutated proteins or molecular circuitry essential to cancer cells. The complexity of cell regulation and the profound cellular reprogramming associated with malignant transformation, however, provide formidable barriers to elucidating disease-critical genetic circuitry. We propose that linking genetic perturbation experiments with statistical modeling can provide inroads to discovery of functionally relevant gene regulatory network (GRN) architecture and thus valuable prediction and validation of genetic interactions in cancer gene networks, which ultimately may inform targeting of next generation cancer interventions.

Gene regulatory network (GRN) modeling seeks to infer dependencies between genes and thereby provide insight into the regulatory relationships that exist within a cell. Early methods to estimate GRNs focused primarily on data arising from targeted experimental perturbations^1, 2^; however, these methods relied on simplifying assumptions to reduce the computational demands of these methods. As the size and scope of publicly available transcriptomic data rapidly increased due to the formation of large cancer consortia such as The Cancer Genome Atlas (TCGA), GRN modeling methodology transitioned to leveraging observational data, which rely on correlation structure in the data produced by natural biological covariation to infer regulatory relationships^3, 4, 5, 6, 7^. While application of these methods have provided insights into the regulatory systems crucial to malignancy, these modeling approaches are hindered by technical sources of covariation, such as batch effects or variable sample composition, either masking or mimicking true biological dependencies^8^. Additionally, modeling of GRNs remains inherently difficult due to high order non-linear interactions and unmeasured sources of biological variability that influence the GRN.

Methodological and computational advances have greatly aided our ability to model GRNs, which in turn has advanced our understanding of the processes involved in both normal cellular function and disease progression. While much effort has been dedicated to estimate GRNs from observational data, experimental perturbations allow direct measurement of the effect that a change in one gene has on other measured genes, which greatly facilitates estimation of GRNs^9, 10, 11^. Recent advances in computational hardware and parallelization provide an opportunity to revisit the potential of perturbation based GRN estimation without the need to impose biologically implausible restrictions on these models.

In this paper, we propose a GRN modeling procedure, termed TopNet, based on underlying gene perturbation data that addresses the core challenges of GRN estimation: preprocessing and analysis of noisy experimental data, quantification of uncertainty in gene expression changes following perturbation, exploration of a vast model space to find networks supported by the gene perturbation data, and identification of robust, biologically relevant network features. Our approach adopts the ternary network modeling formalism proposed in Almudevar et al ^12^. Specifically, we consider a ternary network modeling procedure that samples networks for which the network attractors agree with the experimental data. In this paper, we propose improvements in both the search algorithm and the manner in which agreement between the attractors and observed data is quantified.

Previously we have been able to identify non-mutant mediator genes critical to the cancer phenotype downstream of oncogenic mutations^13, 14^. Our approach to find such mediators was based on the concept that the profound and multi-faceted genetic reprogramming associated with malignant cell transformation requires cooperation of multiple oncogenic mutations^15, 16^, and that this transition is driven by synergistic modulation of downstream mediators^17, 18, 19^. This class of functionally diverse and apparently unconnected genes was termed ‘cooperation response genes’ (CRGs). Functional analysis of CRGs revealed that >50% of CRGs tested play an essential role for the cancer phenotype^13^, and that specific CRGs represented newly identified, cancer cell-specific vulnerabilities^20, 21, 22^. Given the functional diversity of CRGs and the robust and redundant nature of regulatory circuitry in mammalian cells, this high frequency of impact on the cancer phenotype by individual CRG perturbations is contrary to expectation, unless CRGs act in a concerted manner. Our work demonstrates that CRGs act within a strong network of unexpected genetic interdependencies that appears critical to the robustness of the malignant phenotype.

## Results

### CRGs contribute to tumor formation capacity

Genetic perturbation of individual CRGs in cells transformed by p53^175H^ and Ras^V12^ (mp53/Ras cells), aimed to re-establish levels of gene expression found in non-transformed parental young adult mouse colon (YAMC) cells, results in a reduction of tumorigenicity in the majority of cases^13^. Extensive analyses presented here recapitulate our earlier results. We now show an entirely new and much larger set of 49 CRG perturbations (Figure 1 and Supplemental Figures 1 – 2). Of these, 26 perturbations resulted in significantly reduced tumor formation capacity of the target cells when implanted into immune compromised mice. Together with our previous work, these data demonstrate that 41 out of 75 CRGs tested (55%) are critical to the cancer phenotype (Figure 1A).

**Figure 1:**
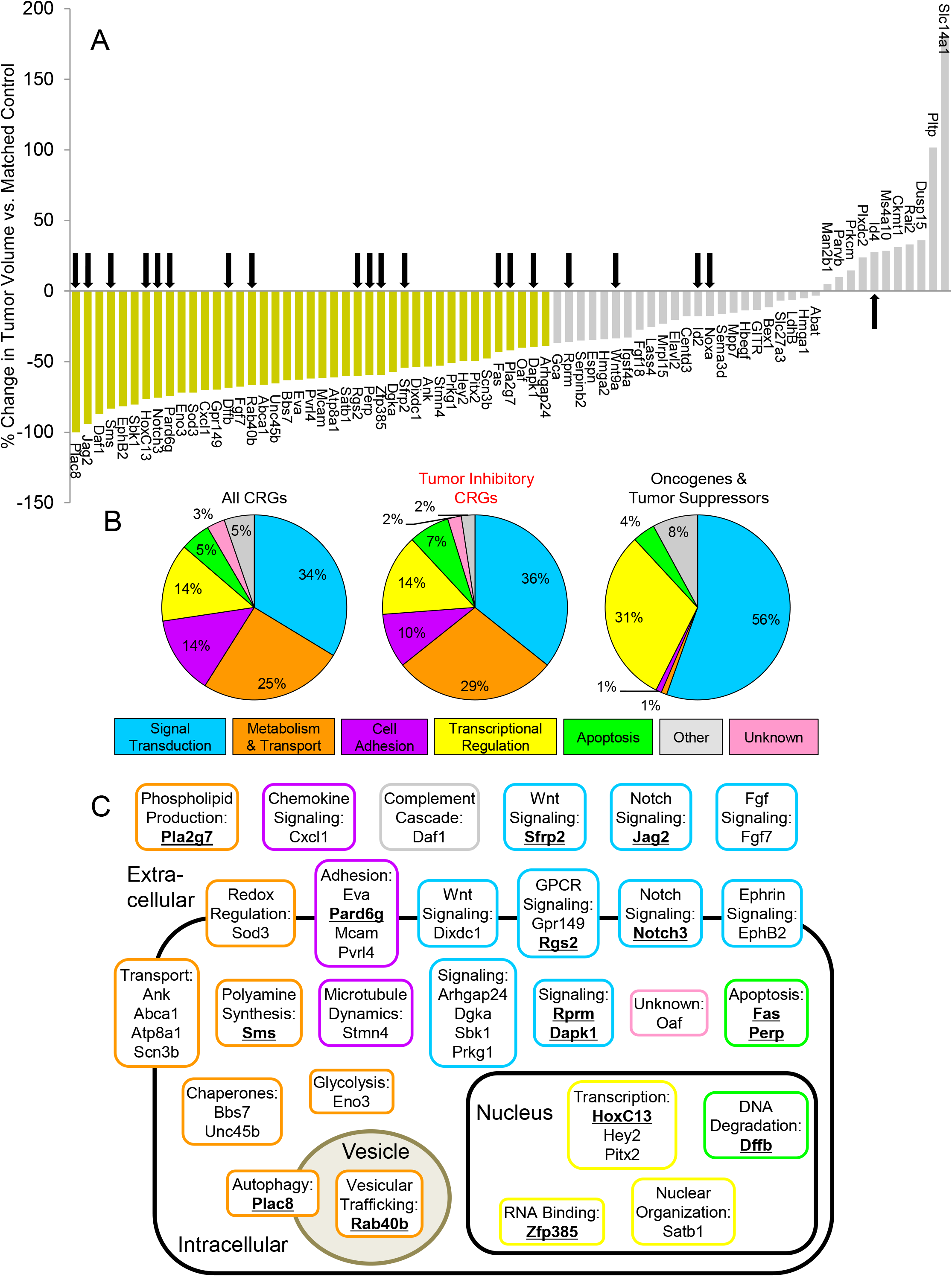
Impact of CRG perturbations on tumor formation. (A) Waterfall plot indicates percent change in endpoint tumor volume of allografts following perturbation of individual CRGs in mp53/Ras cells. Perturbations significantly decreasing tumor size, as compared to matched controls are shown in gold (p<0.05, unadj. Wilcoxon signed-rank test). Those with no significant reduction in tumor size are shown in gray. Arrows indicate genes chosen for inclusion in gene regulatory network modeling. (B) Pie charts indicate the proportions of functional gene annotations according to the Gene Ontology database for all CRGs, tumor-inhibitory CRGs and classical oncogenes and tumor suppressors, respectively. Colors signify biological processes, as indicated. (C) Scheme summarizes the cellular localization and cell biological functions of proteins encoded by CRGs, colored as in (B). CRGs chosen for inclusion in gene regulatory network analysis are indicated by bold, underlined text.

CRGs regulate diverse hallmarks of the malignant phenotype. The distribution of associated cell processes among those genes with a demonstrated role in tumor regulation is highly similar to the distribution across all CRGs but distinct from known oncogenes and tumor suppressors (Figure 1B). The genes whose perturbation impacts tumor formation vary in their localization from nuclear to intracellular to extracellular function and are associated with numerous regulatory pathways (Figure 1C). Notably, many of the CRGs that are implicated in control of tumor formation capacity are associated with processes such as metabolism and cell adhesion, regulators of which are not detected among frequently mutated oncogenes and tumor suppressor genes. Thus, we consider the CRGs as critical mediators of oncogene cooperation that transmit and carry out the effects of those oncogenic mutations.

### Individual CRG perturbations impact expression of other CRGs

CRGs function in many diverse pathways and mechanisms (Figure 1B, C), and interrelationships between most of these genes have not been previously reported. In contrast, the high frequency by which CRG perturbations impact the cancer phenotype and the manner in which the CRGs themselves were identified, by virtue of their synergistic response to multiple oncogenic mutations, led us to predict that CRGs function within a regulatory gene network. We thus investigated the extent of mutual control of gene expression among 20 CRGs, which were selected to represent the spectrum of functional classes and tumor inhibitory effects observed in the full CRG set (indicated in Figure 1A, 1C). Using a set of mp53/Ras cell populations, each harboring one of twenty individual CRG perturbations, we compared mRNA expression profiles of perturbed cells with corresponding controls to detect changes in expression of the selected CRGs. These perturbations varied in their effects on tumor growth and were chosen from genes up- or down-regulated in transformed mp53/Ras cells as compared to the parental YAMC cells (Figure 2A). Expression profiles for all 20 CRGs were measured for multiple independent replicates of each CRG perturbation, revealing wide-spread changes in gene expression in the CRG cohort upon perturbation of individual CRGs (Figure 2B).

**Figure 2:**
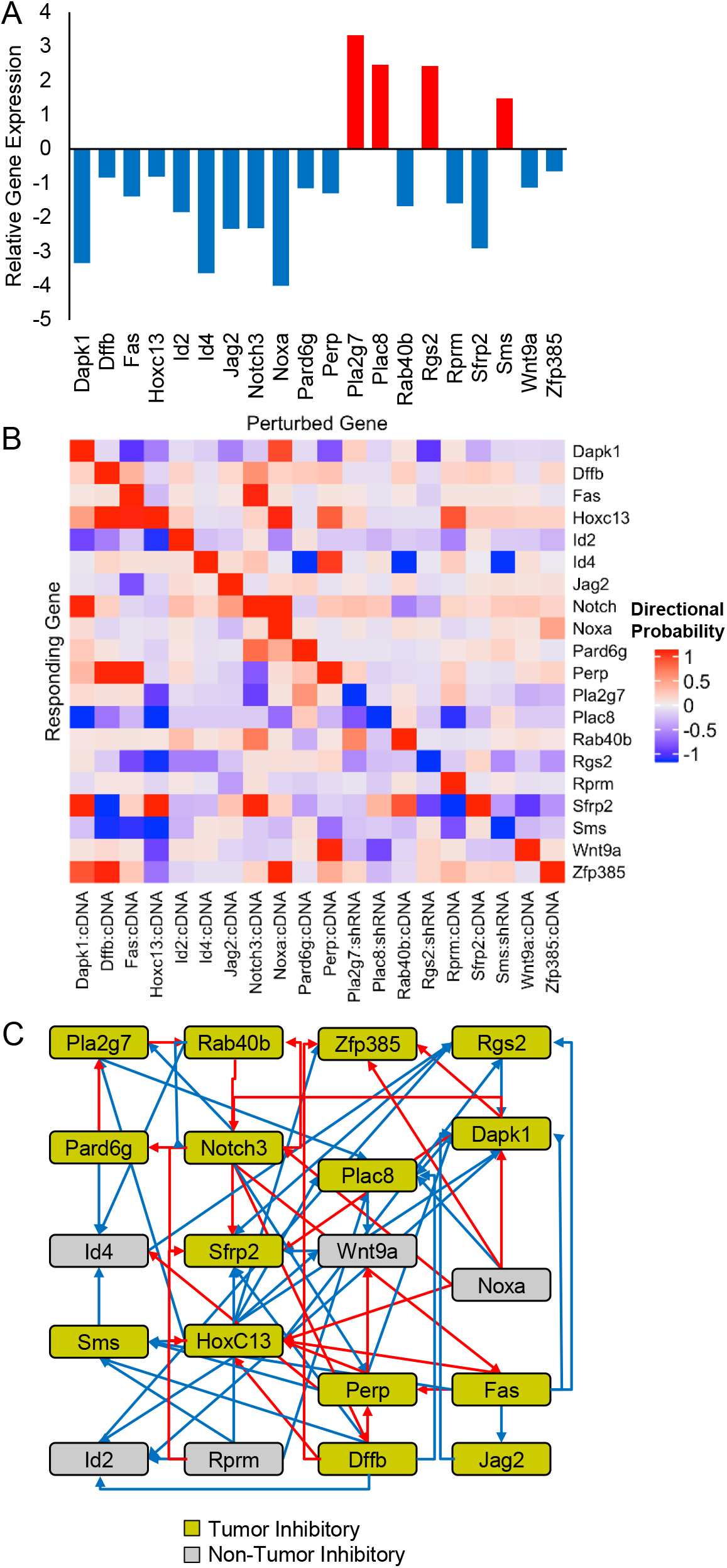
CRG connectivity. (A) Bar graph showing the expression levels of each CRG selected for network analysis, as compared to their baseline expression in non-transformed parental YAMC cells. Genes up-regulated in mp53/Ras vs. YAMC are shown in red, while down-regulated CRGs are shown in blue. Measurements were made by TLDA for multiple replicates of each gene perturbation. (B) Probability of change in expression of each CRG in response to individual CRG perturbations indicated, computed based on mRNA levels detected on TLDA for multiple replicates of each gene perturbation. Red denotes a likely increase in expression of the downstream gene, blue denotes a likely decrease in expression. Note that the diagonal indicates the direction of perturbation, intended to restore levels of gene expression found in parental YAMC cells for each perturbed gene. (C) Connectivity graph showing the influence of given CRGs on the expression of other CRGs. Arrows indicate parent-child connections with |probability| > 0.5, colored as in part A to indicate likely direction of change in the child. Nodes are colored as in Figure 1 to indicate tumor inhibitory effects of perturbation of the indicated CRGs (gold = inhibitory, gray = no significant effect). Placement of nodes was chosen for comparison to Figure 5.

The high degree of connectivity between the selected CRGs is shown in Figure 2C and Supplemental Figure 3A. Key parameters that describe the interconnectedness of genes in such a graph are the *out-degree*, i.e. the number of children of a given gene, and the *in-degree*, the number of parents of a given gene. Of the 20 perturbations, eight produced changes in the expression of 5 or more of the other CRGs, with one perturbation, restoration of HoxC13 expression, having the highest out-degree, i.e. affecting the expression of 9 other CRGs (Supplemental Figure 3A and B). Only two perturbations had an out-degree of zero, indicating that they did not produce a change in any other CRG. Moreover, there was substantial variability in the in-degree among these CRGs, ranging from two CRGs not responding to any other perturbations, to the one CRG, Sfrp2, with the highest in-degree, responding to perturbation of 8 other CRGs (Supplemental Figure 3A and C). Taken together these results highlight the remarkably high connectedness of these 20 CRGs. Given the numerous interactions between this subset of CRGs, we employed statistical modeling to understand the flow of information in this gene regulatory network and associated network architecture.

**Figure 3:**
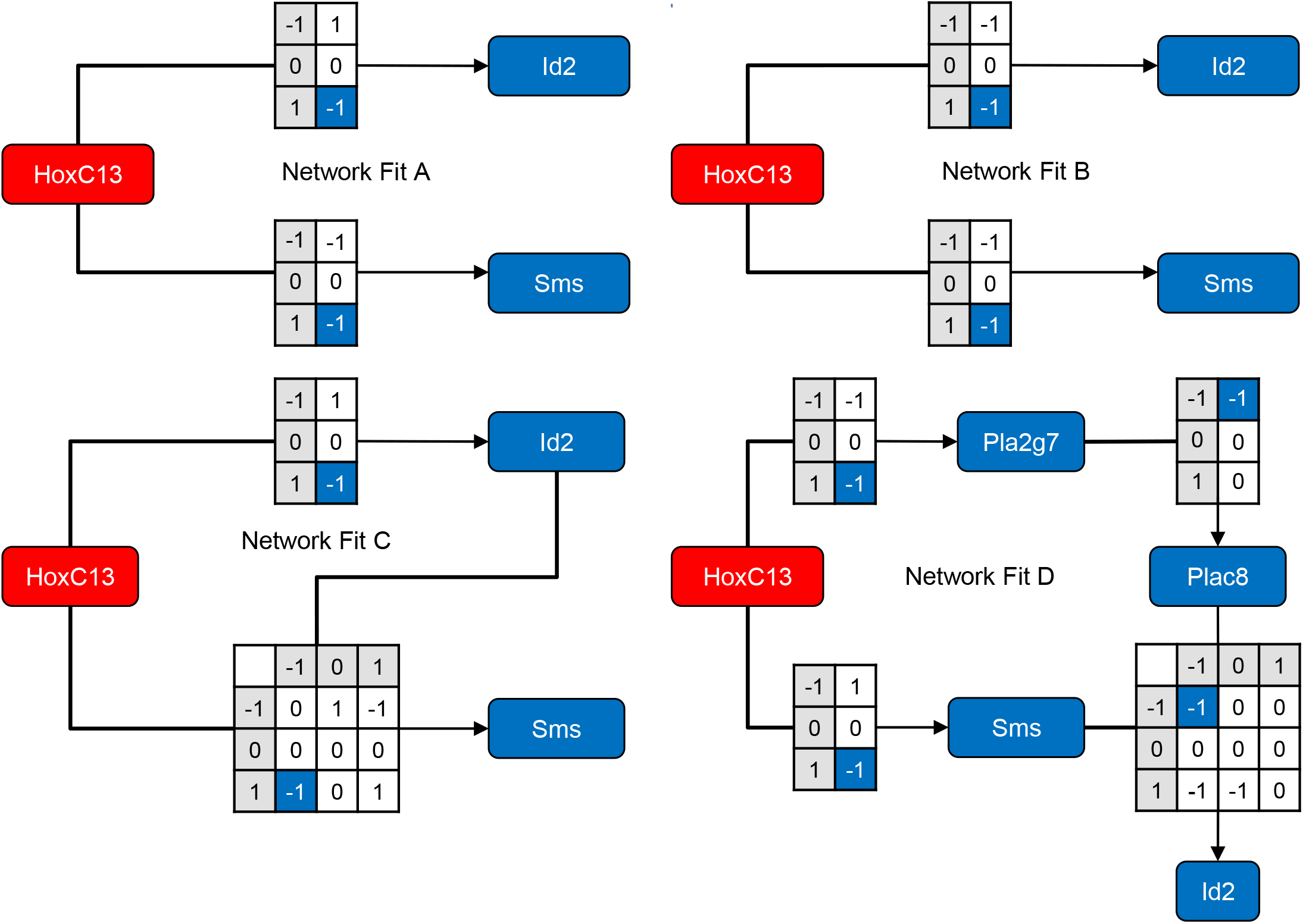
Transition functions encode network behavior. Paths from perturbation of parent node HoxC13 to its possible children are diagrammed. Four network fits of 100 computed are represented. Marginal effects (single parent) and joint effects (multiple parents) are shown with transition function tables. For simplicity of visualization, the maximum in-degree shown here is two but the models applied allow for up to four parents to influence a given child node’s expression. Red nodes indicate that the gene increases its expression, while blue denotes reduced expression.

### TopNet reveals rules governing attractor states

To identify gene interactions relevant to the malignant phenotype, we developed TopNet, a ternary network modeling approach based on perturbations capable of up- or down-regulation of gene expression that accounts for uncertainty in the underlying gene perturbation data. This appeared suitable, as CRGs comprise both up- and down-regulated genes, including genes with very low expression levels^13^.

Estimation of a GRN typically focuses on inferring individual interactions (edges) between genes and/or proteins (nodes). The collective behavior of these interactions is then studied as an emergent property of the low-level network architecture. This is a natural strategy when inferring GRNs based on measurement of pair-wise interactions – e.g. transcription factor binding (either via ChIP-seq or promoter sequence motifs) or protein-protein interaction data. However, gene perturbation experiments remain the most reliable approach to investigate GRNs and to predict cellular response to interventions^9, 10, 11^. The gene expression changes in response to perturbation provide information not on the direct interactions between genes but rather on the altered steady-state expression produced by the perturbation. In other words, the estimation procedure can be framed as a search through the vast space of network models for those that produce the observed data^12^. In contrast to previous work, TopNet gives more weight to experimental results with higher certainty through application of a finite mixture model that provides probabilities of up-/ down-regulation for each gene in response to a given perturbation (Figure 2B). Incorporating these probabilities into the network-modeling algorithm allows us to more accurately quantify our confidence in the estimated dependences between genes.

To assess potential interdependence between the 20 CRGs selected for network estimation, we applied the TopNet algorithm, which uses a ternary network model that accounts for the dynamic, cyclic, and non-linear nature of GRNs and facilitates the evaluation of uncertainty^12^. Specifically, TopNet assumes that changes in gene expression can be modeled by one of three states – down-regulation (−1), baseline expression (0), or up-regulation (+1) – and that the network is governed by deterministic transition functions that encode the complex regulatory relationships between genes (examples are shown in Figure 3A-D). Discretization captures the majority of information and inherently provides robustness^23, 24, 25, 26^. Additional details of the TopNet modeling algorithm are described in the Methods and Supplemental Files 1-3.

Following any given perturbation, the network will eventually reach a steady state, called an attractor. This attractor may be a new state to which the cell is driven by the action of the gene perturbation, or may represent a return to the baseline state, in this case that of a transformed mp53/Ras cell. Recall that the experimental data represent a measurement of the steady state expression of the cells; therefore, the ability of a network model to represent experimental data can be assessed by comparing these gene expression measurements with the attractors predicted by the network model^12^. Specifically, starting from a null model, TopNet randomly proposes small changes in the network structure and scores the proposed new network based on the similarity between its attractors and the observed steady state data (Figure 4; Methods and Supplemental Files 1-3). This iterative search algorithm can proceed quite rapidly because TopNet can easily compute the attractor for each perturbation and assess the similarity of the resulting attractors with the corresponding observed steady state data. Ultimately, TopNet identifies a network model whose attractors match the observed data well.

**Figure 4:**
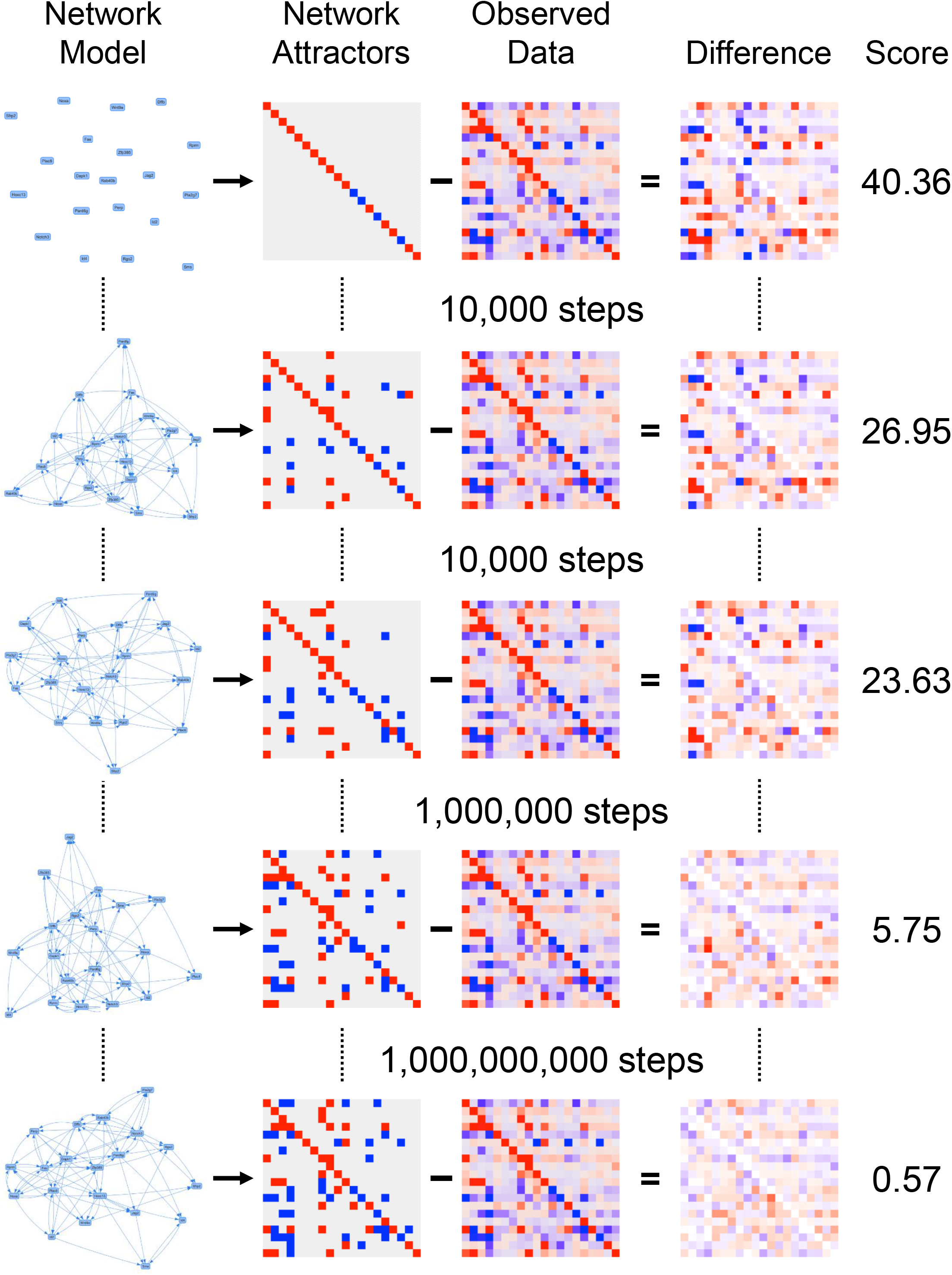
Gene regulatory network reconstruction process. The network modeling algorithm begins with a null network in which no gene affects any other gene (top left). This produces a set of null attractors wherein perturbation of any gene does not produce a change in any other gene. The network models are scored by summing the absolute value of the differences between the probabilities of up- or down-regulation and the attractors, which are then standardized by subtracting the best theoretically possible score. Comparing the null attractors to the calculated probabilities results in substantial differences, which produce a score of 40.36, indicating that the network model does not explain the observed data well. Random changes in topology and/or transition functions are made to the network model and the process is repeated. Here, we show snapshots of the process after the first 10,000 steps, the next 10,000 steps, then after an additional 1,000,000 steps, and finally after 1,000,000,000 steps at which point the network model produces attractors that match the probabilities calculated from the data very well.

### Uncertainty quantification allows the identification of high confidence features of the network topology

There are often many network models that fit the observed data equally well, and the model space is huge -- for the network model presented here, there are approximately 2.48×10^763^ potential networks. To more efficiently explore this vast model space and avoid becoming trapped in local optima, we implemented a replica exchange Monte Carlo algorithm^27^ that allowed us to parallelize the search over the nodes of a high-performance computing cluster. Unlike previous approaches, here we have modeled and addressed non-random missing data^28^, batch effects, and uncertainty in estimates of differential expression (Supplemental File 3). These improvements in data analysis prior to network estimation were crucial to accurately model our data.

Due to the vast number of potential networks, it is exceedingly unlikely that there exists a single “best” network; therefore, as previously proposed in Almudevar et al.^12^, TopNet repeats the estimation procedure many times to produce a sample of networks from the space of models that explain the observed data well. This collection of network fits can then be used to quantify the degree of support for a given aspect of the network by calculating the proportion of networks in which a given feature or features are present. For example, based on the four network models shown in Figure 3, we could state that HoxC13 is likely a direct parent of Sms because that relationship is present in all 4 models, whereas we have less certainty about the relationship between HoxC13 and Id2 because it is only a direct parent in 3 of the 4 models, remembering that the four models shown are only a sub-sample of all that were computed.

### Assessments of model fit and complexity demonstrate the significance of the estimated network

One question of interest is whether it is significant that TopNet can obtain a network model producing attractors that match the observed steady-state data. We examined this by permuting the network input data. For each gene, we permuted its response to all of the experiments. This retains the number of experiments to which each gene responds. In other words, the number of parents in a connectivity graph remains constant but we randomly change the identity of parent genes. We then used TopNet to fit a network model to the permuted data using the same parameters as for the unpermuted data. We repeated this process 250 times and compared the network scores from the real data to the permuted data. The network scores based on the real data are better (i.e., lower) than nearly all the scores based on the permuted data (Supplemental Figure 4, empirical p-value <0.01). Thus, the currently utilized network parameters allow us to obtain good scores for the real data but not for the permuted data. This suggests that we are not extensively overfitting these data and that the current network constraints are reasonable.

We also examined whether we could obtain similar fits with a simpler network model. Specifically, we tested whether reducing the number of parents (in-degree) still allowed us to fit the observed data reasonably well. To investigate this, we re-ran the network modeling algorithm with the *in-degree cap* reduced from 4 to 3 (run 100 times). Reducing the in-degree results in a substantial increase in the network scores (Supplemental Figure 4), lending further support to the choice of a maximum in-degree of four for these data. We also re-ran the network-modeling algorithm with the in-degree cap increased to 5 (run 20 times). This produced network models with scores slightly better than the in-degree 4 models (Supplemental Figure 4). We also compared the scores from the in-degree 3 and 5 fits based on the real data to fits based on permuted data. Regardless of the in-degree cap, better scores were achieved using the real data; however, with an in-degree cap of 5, it was at times possible to fit the permuted data perfectly, suggesting that the in-degree 5 model is too flexible. Furthermore, the separation between real and permuted scores was greatest for an in-degree cap of 4, which lends further support to the choice of a maximum in-degree of four for these data.

### Modeling reveals robust topological features of the CRG regulatory network

The result of the TopNet network modeling procedure is a sample of network models all of which explain the observed data well. Based on this sample of network models, we are able to form inferential statements about specific network features by averaging across the network models. For example, in the case of topological relationships, we calculate the proportion of networks in which a given gene is a parent of another gene. For the CRG network, we report results based on 100 independent network fits. In Figure 5, we show edges present in at least 80% of the network models, which represent regulatory relationships that are strongly supported by the observed data.

**Figure 5:**
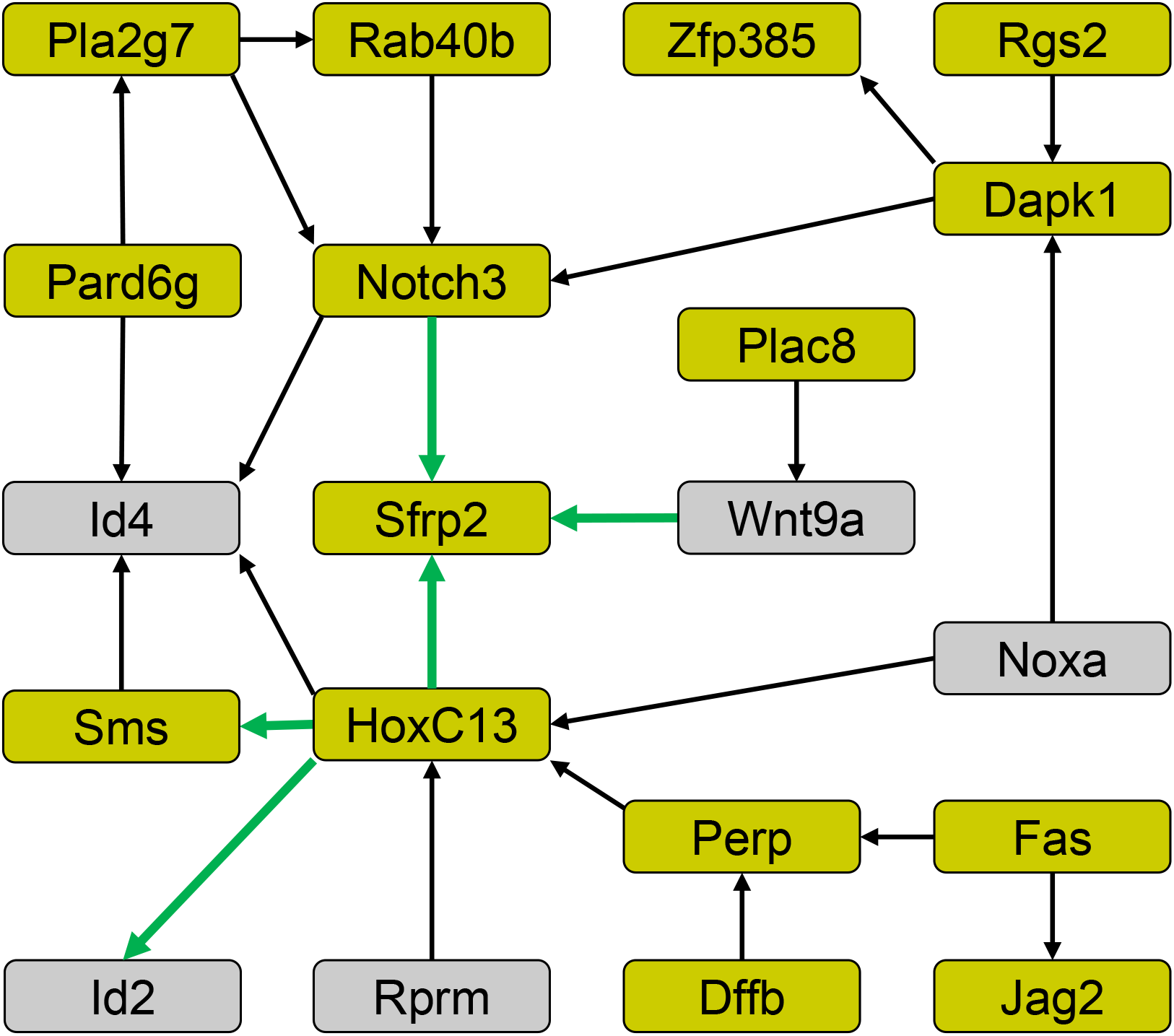
Summary of CRG network topology. The network models produced by 100 independent fits are mapped by reporting the proportion of networks in which an edge (parent – child relationship) appears in more than 80% of the network models. Node color indicates the effect of perturbation of that node on tumor formation capacity (as shown in Figure 1 and used in Figure 2, gold = tumor inhibitory perturbation, gray = no significant effect). Tested epistatic interactions are indicated by green arrows.

Modeling gene regulatory relationships between CRGs allows us to make and test predictions about the role such relationships might play in maintaining or generating a given biological state; here the malignant state. Using the network topology (Figure 5) and information from the underlying transition functions that describe the specific interdependencies between network nodes, we set out to test the impact of two central players common to all the network models, HoxC13 and Sfrp2, on which we focused further experiments.

### Epistatic interactions between functionally diverse mediator genes

First, we tested whether the predicted parent-child relationships between CRGs relate to biological dependence between upstream and downstream genes. In order to do this, we identified parent-child pairs where perturbation of the upstream gene has a reproducible biological effect, in this case inhibition of tumor growth *in vivo*. We focused on the parent node with the most children, HoxC13, a homeobox transcription factor^13, 29^, perturbation of which produced some of the strongest anti-tumor effects observed in our experiments. For double perturbations, we selected the parent-child relationships between HoxC13 and its predicted

children: Sms, the gene encoding spermine synthase30, Id2, a gene encoding a member of the helix-loop-helix only and inhibitor of DNA binding protein family^31^, and Sfrp2, encoding a member of the family of secreted Frizzled-related proteins, antagonists of Wnt signaling^32^ (Figure 6A). Note that the relationship between HoxC13 and Id4, while predicted by the network topology, is not observed in the connectivity graph, and thus an appropriate genetic perturbation to test this relationship remains unclear.

**Figure 6:**
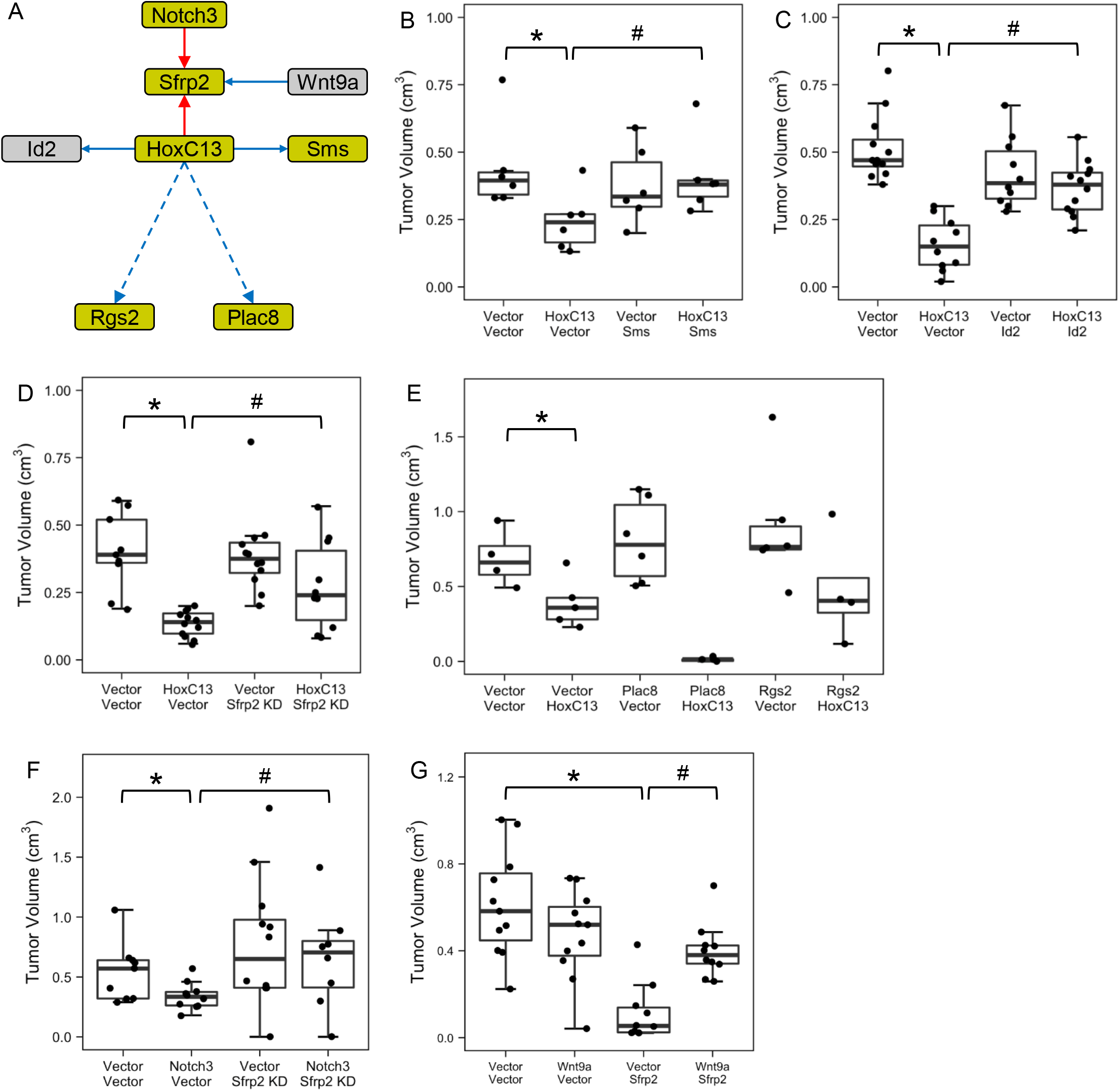
Epistatic CRG interactions impacting cancer phenotype. (A) Diagram of a subnetwork of the GRN shown in Figure 5, whose edges are indicated in solid lines, with the addition of two edges seen only in the connectivity graph but not in the topological map (dashed lines). Red arrows indicate that perturbation of the parent gene increases the child’s expression, while blue arrows denote reduced expression of the child in response to the parent perturbation. (B-G) Box plots show tumor growth in response to indicated single and double perturbations targeting parent node HoxC13 and its predicted children, or child node Sfrp2 and its parents. Perturbations significantly decreasing tumor size, as compared to matched controls are indicated by an asterisk (*, p<0.05, unadj. Wilcoxon signed-rank test). Double gene perturbations showing significantly different tumor size as compared to individual gene perturbations are indicated by a hash mark (#, p<0.1, unadj. Wilcoxon signed-rank test vs. each individual perturbation).

Our experiments testing relationships between HoxC13 and Sms, Id2 or Sfrp2 demonstrate that changes to the expression of these child nodes are required for the biological effects of the parent perturbation (Figure 6B-D, Supplemental Figure 5). Specifically, we find that the ability of HoxC13 to suppress the malignant state is curtailed when either low Sfrp2 levels or high Sms or Id2 levels are enforced. Notably, all three of these genetic interactions represent previously unknown dependencies critical to the cancer phenotype.

The GRN presented makes specific and strong predictions about interactions that may help to maintain the robustness of the network and ultimately the malignant state of the cells. However, parent-child relationships beyond those identified by modeling the GRN can be seen by direct examination of the results of a perturbation on the other genes in consideration (Figure 2, Figure 6A). Specific to HoxC13 as a parent, gene expression profiling reveals that expression of Plac8, encoding a lysosomal protein critical for autophagosome-lysosome fusion in cancer cells^21, 33^, and Rgs2, which encodes a family member of regulators of G protein coupled receptors^34^, are each decreased by HoxC13 perturbation. However, these interactions are not necessary for the GRN to explain the steady state observations of expression of all the genes in the network and thus are not frequently observed in the topological mapping of the GRN. To test the relative importance of interactions identified by network modeling versus those observed by gene expression profiling, we generated double perturbations re-expressing HoxC13 together with either Plac8 or Rgs2. Remarkably, neither of these combinations revealed dependence of the anti-tumor effects of HoxC13 on Plac8 or Rgs2 (Figure 6E, Supplemental Figure 5), although enforcing both HoxC13 and Plac8 expression had an enhanced anti-tumor effect. Together, these results indicate that modeling of the GRN reveals biologically relevant epistatic gene interactions (Figure 5), while simple connectivity graphs (Figure 2B) do not allow for such prioritization.

Last, we examined whether relationships between Sfrp2, the gene affected by the largest proportion of perturbations (Figure 2C, Supplemental Figure 3A), and its predicted parents, HoxC13, Notch3, and Wnt9a (Figures 5 and 6A) are critical to the malignant phenotype. As described above, re-expression of HoxC13 does not significantly inhibit tumor growth when low Sfrp2 levels are maintained. Similarly, the tumor inhibitory effect of Notch3 expression is suppressed when Sfrp2 expression is maintained at low levels (Figure 6F, Supplemental Figure 5). Unlike re-expression of HoxC13, Notch3 or Sfrp2, re-expression of Wnt9a has little to no effect on tumor size (Figure 1A and 6G), while this decreases expression of Sfrp2 (Figure 2A). When Sfrp2 and Wnt9a expression are both increased, we observed a modest but significant reduction in tumor growth compared to controls (p = 0.029, unadj. Wilcoxon signed-rank test) but increased tumor growth compared to Sfrp2 expression alone (Figure 6G, Supplemental Figure 5), consistent with the idea that Sfrp2 and Wnt antagonize each other both at the level of gene expression and Wnt signaling activity^32^.

Taken together, here we demonstrate that CRGs function within a gene regulatory network that contributes to the malignant phenotype. Our results reveal that numerous tumor regulatory interactions are detected via attractor-based ternary GRN modeling through the TopNet algorithm. Notably, high confidence topological features of the network model are frequently found to be biologically relevant, and thus the network model effectively predicts previously unknown epistatic gene interactions critical to the cancer phenotype.

## Discussion

Here we show that apparently unrelated mediators of the cancer phenotype are linked by a strong network of genetic interactions and thus maintain robustness of the malignant state. Critical to this discovery, we developed and applied TopNet, a novel approach to GRN estimation, capable of identifying a biologically relevant topological model of highly complex genetic interdependencies. TopNet, can accurately model cellular responses to perturbations, produce testable hypotheses regarding non-linear multi-gene interactions, and identify control points in the network architecture. This involves searching through a massive number of potential network models to identify those instances consistent with the observed data. By combining the information from many network models, TopNet prioritizes aspects of the GRN that are most strongly supported by the experimental data.

TopNet is capable of pinpointing key architectural features of cancer cells. Application of TopNet to genetic perturbation experiments involving 20 CRGs has identified high confidence multi-gene interactions involved in maintenance of the cancer phenotype. Further experimentation confirmed all of the tested interactions identified by TopNet as previously unknown epistatic gene interactions, while connectivity alone was not a predictor of epistasis. In fact, all of the relationships between HoxC13 and its children Sms, Sfrp2, or Id2, reported here, were experimentally confirmed to be contributing to the tumor-inhibitory activity of HoxC13, which is downregulated in transformed cells as compared to non-transformed parental controls^13^. Our data suggest that HoxC13 inhibits the pathways downstream of Sms and Sfrp2, i.e. polyamine metabolism and Wnt signaling, and Id2 function, all of which is consistent with our observation of reduced tumor growth capacity of transformed cells in which HoxC13 expression has been elevated to resemble levels observed in non-transformed parental cells. Similarly, we demonstrated functional relevance for all predicted interactions between Sfrp2 and its other high-confidence parents, Notch3 and Wnt9A, indicating that the tumor-inhibitory activity of Notch3 is dependent on Sfrp2, and that Wnt9A can down-regulate Sfrp2 expression and thus neutralize Sfrp2-mediated tumor-inhibition, consistent with activation of Wnt signaling.

In addition to identifying high-confidence dependencies via examination of the network topology, TopNet provides the corresponding transition functions that determine the nature of dependencies between parent and child nodes (e.g. Figure 3). However, one should not assume that either the summarized network topology or the transition functions necessarily represent direct mechanistic relationships in the cell. In fact, the functional diversity of the 20 CRGs examined in this work suggests that most of the identified interactions and dependencies act through unmeasured intermediaries. Nevertheless, armed with information on critical interactions in the cell, one can then dedicate effort to understanding such interactions at a mechanistic level.

A major strength of our work is the use of data from gene perturbation experiments as the input for TopNet modeling. While a number of network modeling approaches have been proposed to identify and understand epistatic interactions in cancer cells, many of these modeling approaches are hindered by being observational in nature, as they rely on correlation structure in the data produced by natural biological covariation^35^. In contrast, network modeling based on perturbation experiments allows one to directly assess cause-effect relationships before creating epistatic maps. This has led to increased interest in perturbation experiments as a means of modeling biological systems, such as the use of Perturb-seq^36, 37, 38, 39^ to measure the effect of perturbations at single-cell resolution.

While high-throughput perturbation approaches, such as Perturb-seq, are currently limited by their reliance on gene extinction as the means for gene perturbation, as well as the reduced sensitivity of single cell RNA sequencing technology, we are intrigued by the possibility of adapting our GRN estimation methodology to larger and potentially more complex networks. Extension of TopNet to such data would allow one to move beyond associative analyses and identify non-linear gene interactions underlying cancer cell robustness. Several computational and statistical challenges will need to be addressed in this context, such as the increased feature space, decreased sensitivity, and unidirectional perturbations. In the near term, targeted Perturb-seq^39^, in which some of these challenges are lessened, is ideally suited for initial methods development and testing. Our general modeling approach has the potential to extract rich biological interactions from these data and open new avenues of biological inquiry.

The complexity and strength of the CRG network revealed by TopNet are both remarkable and unexpected. CRGs are comprised of a diverse set of genes functioning in a multitude of diverging pathways involved in regulating cell signaling, metabolism, gene expression, adhesion and survival. In addition, biologically relevant genetic interactions between any of the 20 CRGs investigated here, to our knowledge, have not been reported. Our observations thus suggest a fundamental role for genetic networks in supporting robustness of the malignant state and reveal network stability per se as a target for cancer intervention. One could imagine that such an approach would be advantageous in the context of emerging treatment resistance. In addition, examination of gene regulatory network architecture also has the potential to uncover a wealth of cancer cell vulnerabilities controlled by non-mutated genes, and ultimately identify druggable targets to guide novel intervention strategies. For example, targeting a metabolic enzyme, such as Sms could serve as a convenient alternative to intervention at the level of HoxC13, as knock-down of Sms expression is similarly effective in tumor inhibition as HoxC13 reactivation. Epistasis analysis, as shown here, thus can effectively be utilized for identification of suitable targets accessible to pharmacological intervention in a largely non-druggable GRN environment.

## Methods

### Cells

Young adult mouse colon (YAMC) cells and derivation of transformed cells with multiple oncogenic lesions were derived and are used as described elsewhere^13, 40^. Briefly, polyclonal cell populations harboring mutant forms of p53 (p53^175H^) and Ha-Ras (Ras^V12^) (abbreviated as mp53/Ras) were derived by retroviral infection of low-passage polyclonal YAMC cells. YAMC cells were cultured on Collagen IV-coated dishes (1 μg/cm^2^ for 1.5 hr at room temp; Sigma) in RPMI 1640 medium (Invitrogen) containing 10% (v/v) fetal bovine serum (FBS) (Hyclone), 1×ITS-A (Invitrogen), 2.5 μg/ml gentamycin (Invitrogen), and 5 U/ml interferon γ (R&D Systems). All experiments testing the effects of genetic perturbations were carried out at the non-permissive temperature for large T function (39°C) and in the absence of interferon γ.

### Genetic Perturbation of Gene Expression

cDNAs expressed via pBabe retroviral vectors or pLenti6 lentiviral vectors and shRNA delivered via pSuper-retro retroviral vectors or pLKO lentiviral vectors were used to generate gene perturbations. These were tested by comparison of RNA expression levels in empty vector-infected cells and cells subjected to gene perturbation via SYBR Green qPCR with gene-specific primers.

cDNA clones were obtained from the IMAGE consortium collection, distributed by Open Biosystems, or PCR-cloned from murine cDNA using sequence-specific primers except for murine Jagged2 (Jag2) and Notch3-intracellular domain (Notch3-ICD) (gifts of Dr. L. Milner). All cDNAs were sequence-verified prior to use and were cloned into the retroviral vector pBabe-puro ^41^.

shRNA molecules were designed using an algorithm^42^, available at http://jura.wi.mit.edu/bioc/siRNAext/home.php, for cloning into the pSuper-retro vector^14^ (Oligoengine) or purchased as clones in the pLKO vector. Target sequences for pSuper-retro cloning were synthesized as forward and reverse oligonucleotides (IDT), which were annealed and cloned into the vector. For each up-regulated gene, we identified two or three independent shRNA target sequences yielding at least 50% reduction in gene expression with the goal to guard against off-target effects (Supplemental Figure 2). For this purpose between four and six shRNA targets for each gene were tested. In the case of SerpinB2, only one shRNA target sequence yielded appropriate levels of knock-down, reducing mRNA expression to levels comparable to those in YAMC cells.

Retroviruses were produced following transient transfection of ΦNX-eco cells, while lentiviruses were produced following transient transfection of 293T cells. Infections were carried out in media with 8 μg/mL polybrene. Selection with 5 μg/mL puromycin, and where applicable, 200 μg/mL hygromycin B, was used to generate polyclonal populations of cells stably expressing the indicated cDNAs and/or shRNAs. To test reproducibility of the effects of CRG gene perturbations on tumor formation 2-4 independent replicates of such cell populations were derived (Supplemental Figure 2).

For combined perturbations, cDNA or shRNA for one gene in the pair was sub-cloned into the appropriate pBabe-hygro or pSuper-retro-hygro retroviral vector, allowing for consecutive, independent selection for each gene perturbation introduced. <u>Quantitation of gene perturbation</u>: The efficiency of gene perturbations was tested by comparison of RNA expression levels in empty vector-infected mp53/Ras cells and cells subjected to gene perturbation. Re-expression or knock-down was also compared with the respective levels of RNA expression in YAMC control cells. For collection of RNA, mp53/Ras cells were grown at the 39°C for 2 days, followed by serum withdrawal for 24 hr. Total RNA was extracted from cells following the standard RNeasy Mini Kit protocol for animal cells, with on-column DNase digestion (Qiagen).

SYBR Green-based quantitative PCR was run using cDNA produced as described below for TLDA, with 1x Bio-Rad iQ SYBR Green master mix, 0.2 μM forward and reverse primer mix, with gene-specific qPCR primers for each gene tested. Primers were identified using the Primer Bank database^15^, available at http://pga.mgh.harvard.edu/primerbank/index.html or designed using the IDT PrimerQuest tool (https://www.idtdna.com/Scitools/Applications/Primerquest/). Differential gene expression was calculated by the ΔΔCt method, described below, using RhoA as an endogenous reference gene. Reactions were run on the iCycler (Bio-Rad), as follows: 5 min at 95°C, 45 cycles of 95°C for 30 seconds, 58 to 61°C for 30 seconds, 68 to 72°C for 45 seconds to amplify products, followed by 40 cycles of 94°C with 1°C step-down for 30 seconds to produce melt curves.

As expected, the magnitude of perturbation varies between cDNAs and replicates, and falls into the following groups. For tumor-inhibitory CRGs, all replicates express cDNAs at levels below, at or moderately above YAMC mRNA expression levels with the exception of Pvrl4, for which we cannot exclude the possibility that its tumor inhibitory effects are due to over-expression of the cDNA. For non-tumor-inhibitory CRGs, cDNA expression levels were found at or above the levels of the corresponding YAMC mRNAs (Supplemental Figure 2).

### Allograft Assays

Tumor formation was assessed by sub-cutaneous injection of cells into CD-1 nude mice (Crl: CD-1-Foxn1^nu^, Charles River Laboratories). Murine mp53/Ras cells were grown at 39°C for 2 days prior to injection and implanted via sub-cutaneous injection of 5×10^5^ mp53/Ras in RPMI 1640 with no additives. For each replicate of all gene perturbations, 4 – 8 injections were performed for perturbed cells and vector controls. Tumor size was measured by caliper at 2, 3 and 4 weeks post-injection. Tumor volume was calculated by the formula volume=(4/3)Πr^3^, using the average of two radius measurements. Tumor reduction was calculated based on the average tumor volume following each gene perturbation as compared to the directly matched vector control tumors. Significance of difference in tumor size was calculated by the Wilcoxon signed-rank test using directly matching vector control cells for each perturbation.

### TLDA QPCR

Expression values for each of the CRGs were derived from TaqMan Low-Density QPCR Array (TLDA) data. The TaqMan Low-Density Array (Invitrogen, formerly Applied Biosystems) consists of TaqMan qPCR reactions representing the cooperation response genes and control genes (18S rRNA, GAPDH) in a microfluidic card. TLDA were used to measure gene expression patterns following genetic perturbation of CRGs. To generate cDNA for qPCR analysis, quadruplicate samples of mRNA from mp53/Ras cells harboring CRG perturbations were isolated and 10 µg/sample were mixed with 1x SuperScript II reverse transcriptase buffer, 10 mM DTT, 400 µM dNTP mixture, 0.3 ng random hexamer primer, 2 µL RNaseOUT RNase inhibitor and 2 µL of SuperScript II reverse transcriptase in a 100 µL reaction (all components from Invitrogen). RT reactions were carried out by denaturing RNA at 70°C for 10 minutes, plunging RNA on to ice, adding other components, incubating at 42°C for 1 hour and heat inactivating the RT enzyme by a final incubation at 70°C for 10 minutes.

For each sample, 82 µL of cDNA was combined with 328 µl of nuclease free water (Invitrogen) and an equal volume of TaqMan Universal PCR Master Mix No AmpErase UNG (Applied Biosystems). The mixture was loaded into each of 8 ports on the card at 100 µL per port. Each reaction contained forward and reverse primer at a final concentration of 900 nM and a TaqMan MGB probe (6-FAM) at 250 nM final concentration. The cards were sealed with a TaqMan Low-Density Array Sealer (Applied Biosystems) to prevent cross-contamination. The real-time RT-PCR amplifications were run on an ABI Prism 7900HT Sequence Detection System (Applied Biosystems) with a TaqMan Low Density Array Upgrade. Thermal cycling conditions were as follows: 2 min at 50°C, 10 min at 94.5°C, 40 cycles of 97°C for 30 seconds, and annealing and extension at 59.7°C for 1 minute. Each individual replicate cDNA sample was processed on a separate card.

### TopNet

Our approach to topological network analysis consists of the following specific steps:

1. Non-detect imputation through application of an ECM algorithm to non-random missing data^28^
2. Calculation of z-scores for each perturbation based on ΔΔCt values to standardize expression fold changes.
3. Calculation of the probability of up- or down-regulation in response to each perturbation by fitting a uniform/normal/uniform mixture model to the z-scores calculated in step 2.
4. Identification of ternary network models^12^ which minimize the L1 distance between the network attractors and the estimated probabilities from step 3. For the network model presented in this manuscript there are approximately 2.48×10^763^ potential networks, so we use a novel implementation of replica exchange Monte Carlo^27^ to search the model space in parallel across the nodes of a high-performance computing cluster.
5. Summarization of network topology by reporting parent-child relationships that are present in the vast majority, here at least 80 out of 100, of independent network fits.
6. Generation of testable hypotheses regarding interdependencies among genes in the network via examination of the network topology and the biological effects of perturbation of the genes in the network, in this manuscript inhibition of tumor growth *in vivo*. Several of these steps are described in more detail in the following subsections.

### Preprocessing of Gene Expression Data

Gene expression values were derived using SDS 2.0 and 3.0 software packages (Applied Biosystems). As previously reported in other data sets, we observed a strong dependency between the proportion of non-detects (those reactions failing to produce fluorescence values above a certain threshold) and the average observed expression value. Non-detects were treated as non-random missing data and imputed using the R/Bioconductor package non-detects^28^. The R/Bioconductor HTqPCR package^43^ was used to normalize the data to Becn1 expression, which was shown to have relatively low variability across replicate control samples, and accounts for much of the technical variability between samples (detailed in Supplemental File 3).

Differential gene expression was calculated by the ΔΔCt method. Briefly, using threshold cycle (Ct) for each gene, change in gene expression was calculated for each sample comparison by the formulae:

1. ΔCt_(test sample)_ = Ct_(target gene, test sample)_ – Ct_(reference gene, test sample)_
2. ΔCt_(control sample)_ = Ct_(target gene, control sample)_ – Ct_(reference gene, control sample)_
3. ΔΔCt=ΔCt_(test)_-ΔCt_(calibrator)_

In order to incorporate uncertainty into subsequent analyses, we estimate the probability that a gene is up-/ down-regulated in response to each perturbation by fitting a uniform / normal / uniform mixture model to approximate z-scores for each perturbation. This approach is similar to the probability of expression (POE) algorithm^24^.

### Gene Regulatory Network Reconstruction

The methodology used to estimate the GRN uses the theoretical framework developed in Almudevar et al. (2011)^12^. One novel aspect of TopNet’s approach to GRN modeling is a new method of scoring the networks based on a measure of uncertainty in the differential expression estimates. Specifically, the network models are scored by summing the absolute value of the differences between probabilities of up- or down-regulation and the attractors and subtracting the best theoretically possible score. This allows the network model to give more weight to data points with higher certainty (probabilities closer to 0 or 1). As an added benefit, the ability to produce non-integer network scores eased transitions between network models and significantly decreased computational time. Another novel aspect of TopNet is an improved search algorithm that uses replica exchange Monte Carlo^27^ to parallelize the search across many nodes of a high-performance computing cluster. This allowed us to avoid becoming trapped in local optima and to fit larger more complex networks in substantially less time. Results presented in this manuscript are based on network fits each run for 1,000,000,000 cycles in parallel across 20 processors with temperatures ranging from 0.001 to 1. We have also examined the transition functions, attractors, and trajectories all stored in the fits object, available within Supplemental File 1.

### Code Availability

All statistical analyses can be reproduced using the crgnet R package (Supplemental File 1). The pseudocode for the network modeling algorithm is supplied in Supplemental File 2, while the vignette of the crgnet package is included as Supplemental File 3.

## Acknowledgements

We would like to thank the Wilmot Cancer Institute Genomics Research Center and the Center for Integrated Research Computing at the University of Rochester for technical support. The project described in this publication was supported by the National Human Genome Research Institute of the National Institutes of Health under Award Number HG006853 (to M.N.M.) and the National Cancer Institute of the National Institutes of Health under Award Numbers CA138249 and CA197562 (to H.L.). The content is solely the responsibility of the authors and does not necessarily represent the official views of the National Institutes of Health.

## Author Contributions

H.R.M., H.L. and M.N.M. conceived the study. H.R.M., A.A., L.N., J.A., V.B., and B.S. performed gene perturbation experiments and analyzed data from these. H.S. and M.N.M. developed the TopNet algorithm and software. M.N.M. and H.R.M. performed computational and statistical analyses. H.R.M., A.A., H.L. and M.N.M. interpreted the results and wrote the manuscript.

## Notes

### Competing Interest Statement

The authors have declared no competing interest.

