## Supplemental Figures for "Gene network modeling via TopNet reveals robust epistatic interactions between functionally diverse tumor critical mediator genes"

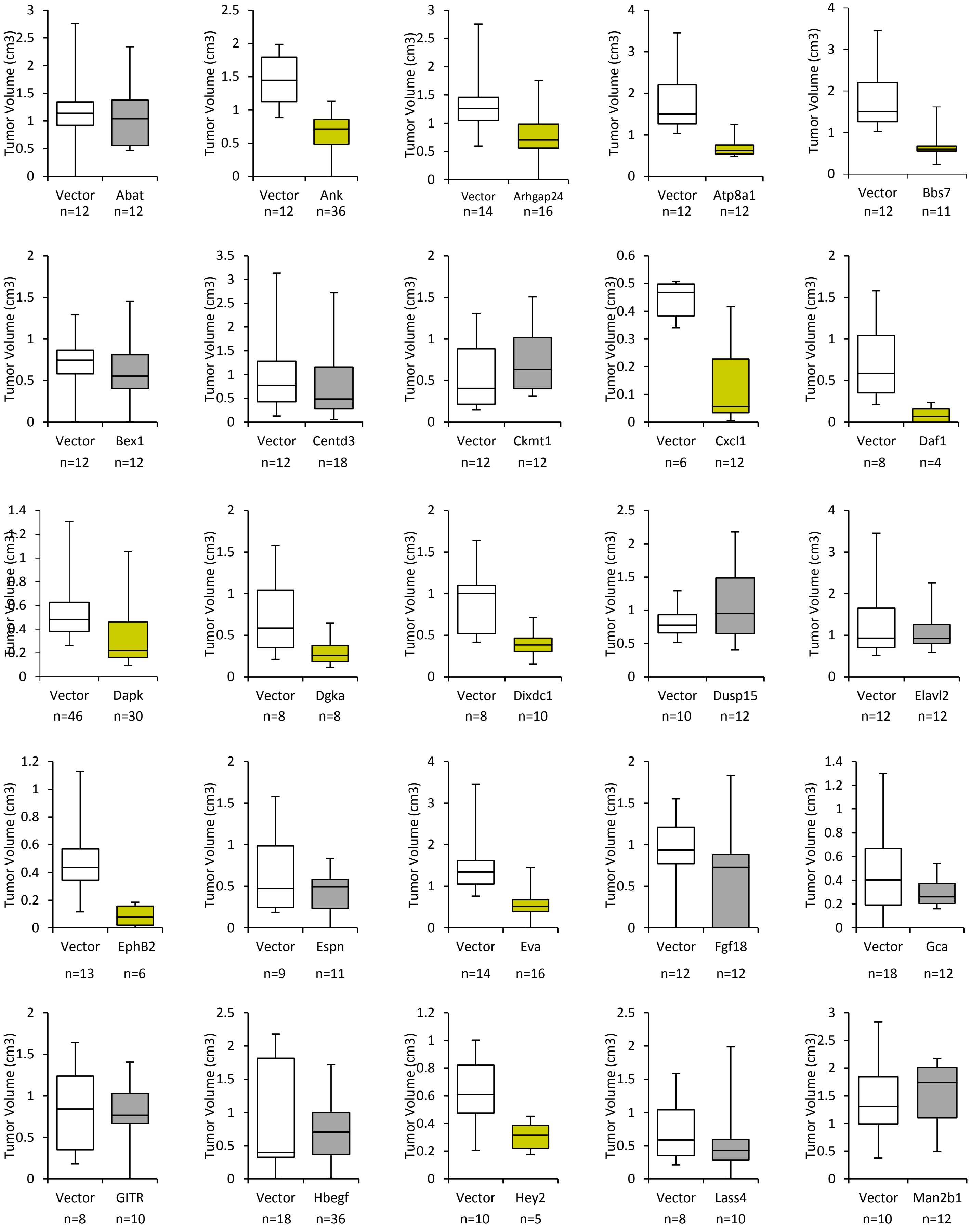


Supplemental Figure 1

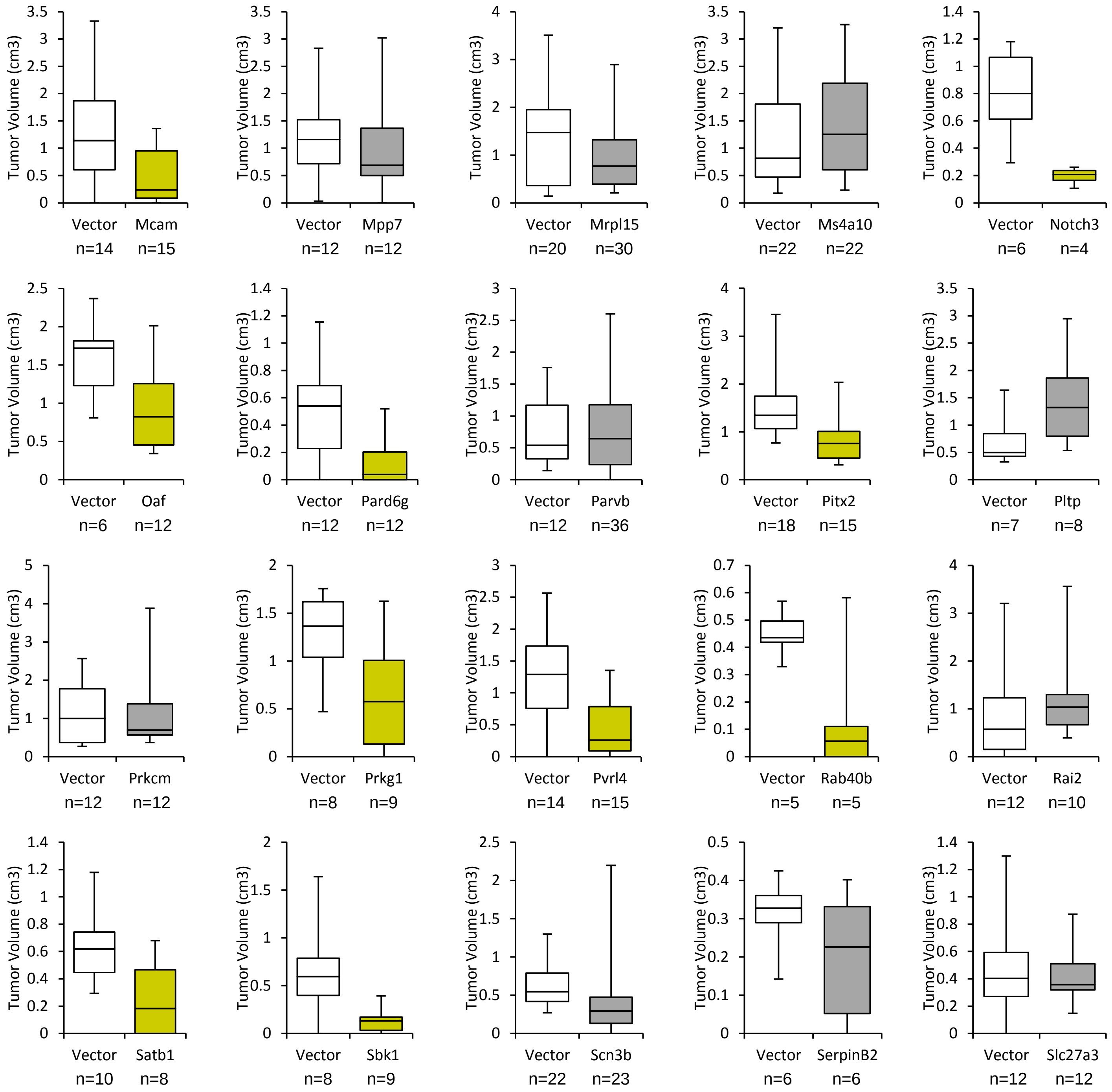

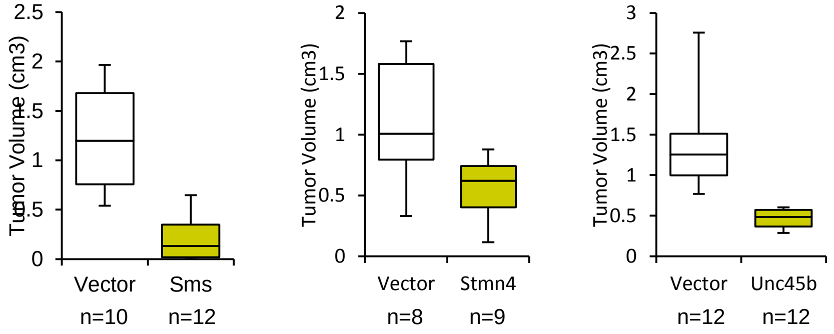


**Supplemental Figure 1: Impact of CRG perturbations on tumor formation of mp53/Ras cells.** Box plots indicate volume (cm3) of tumors formed four weeks after sub-cutaneous injection of cell populations with indicated CRG perturbations (colored boxes), as compared with matched vector controls (white boxes). Boxes indicate range from the first to third quartile of the data, with a line to show median value. Plots are ranked by % change in tumor volume, as in Figure 1. Gold boxes indicate statistically significant change in tumor volume, compared to matched controls (Wilcoxon signed-rank test, unadj. p-value < 0.05). Gray boxes indicate lack of statistical significance. For each perturbation and matched controls, 4-12 implantations were performed per biological replicate for total numbers (n) as indicated.

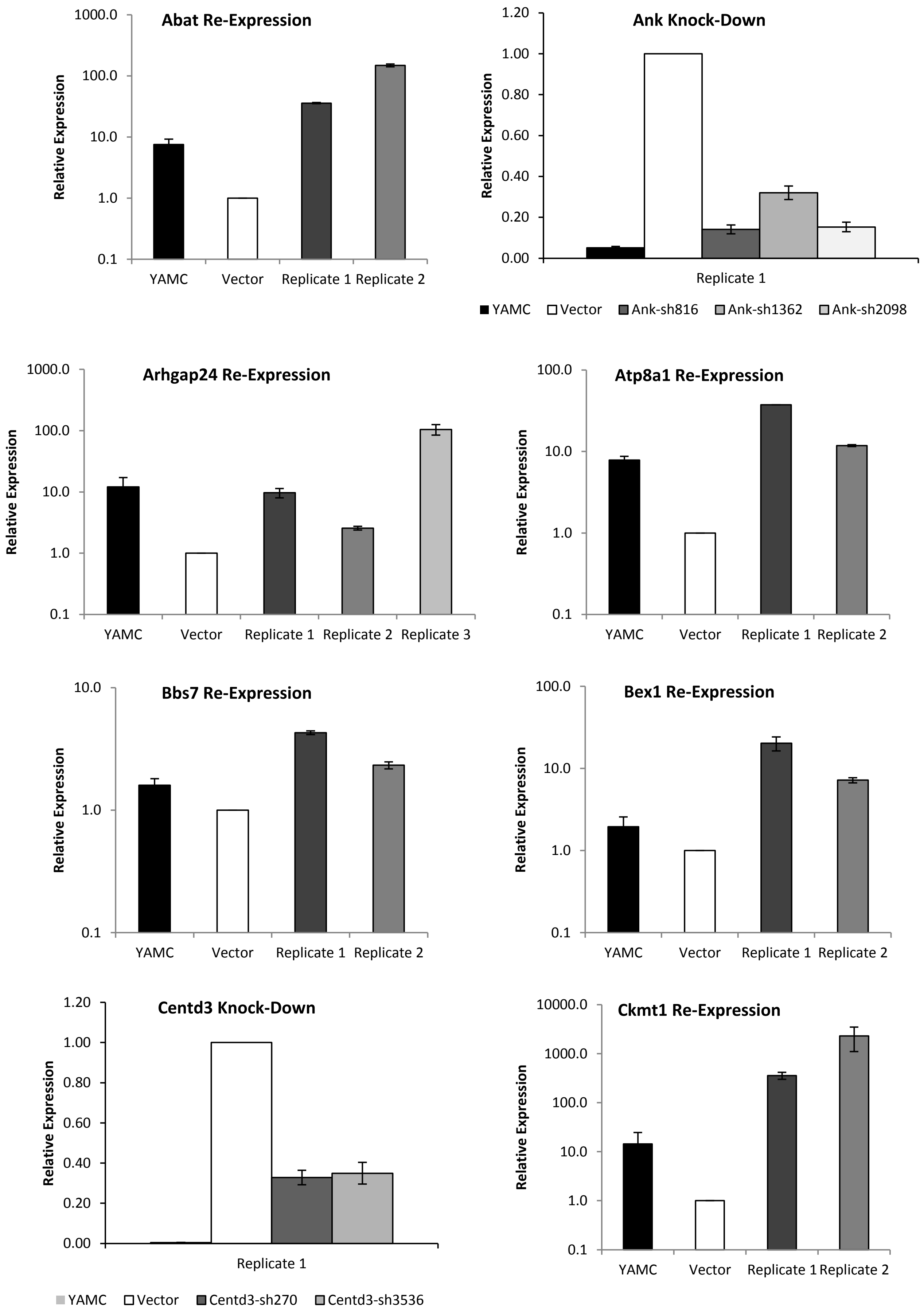


Supplemental Figure 2

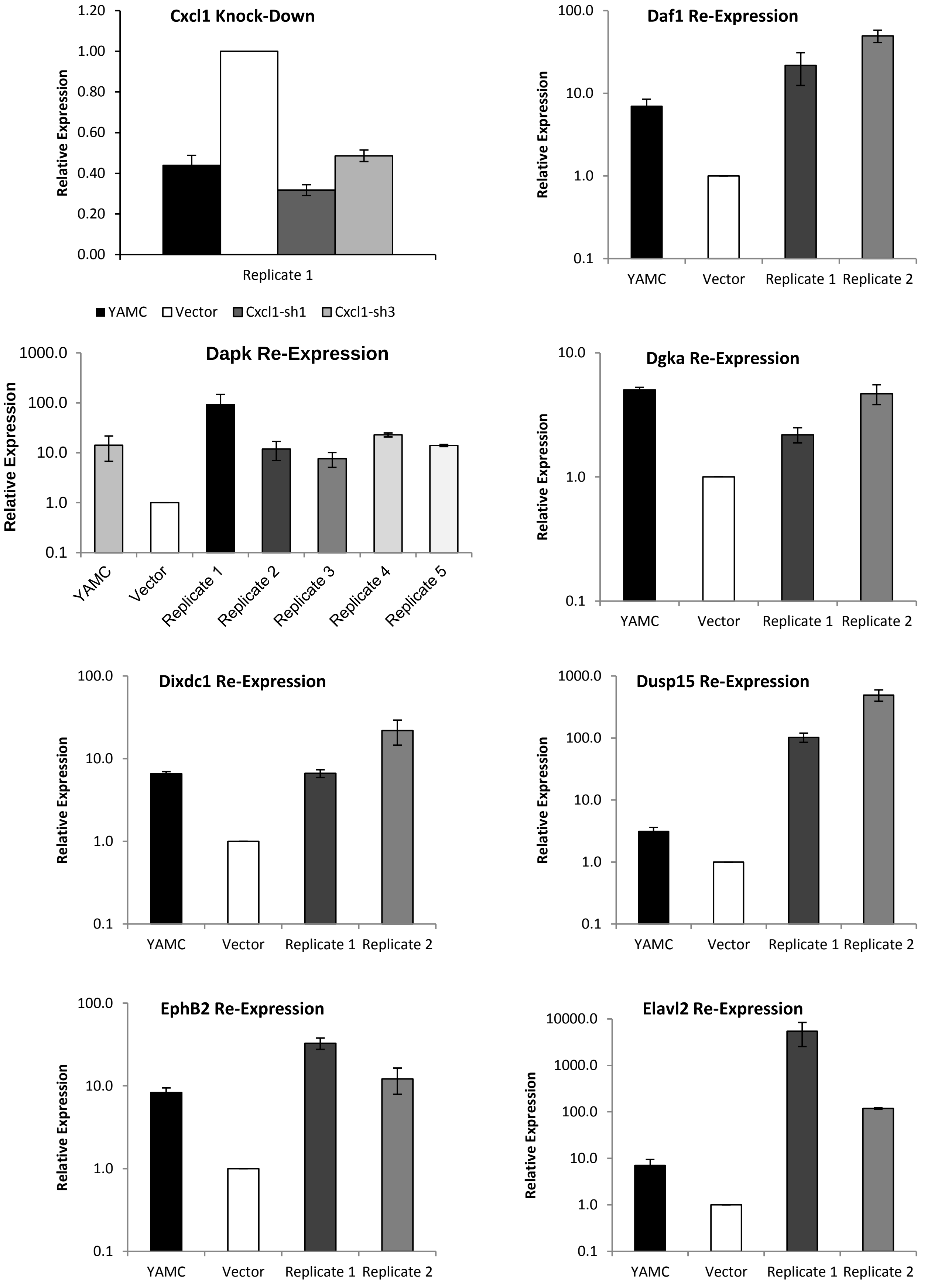


Supplemental Figure 2, continued

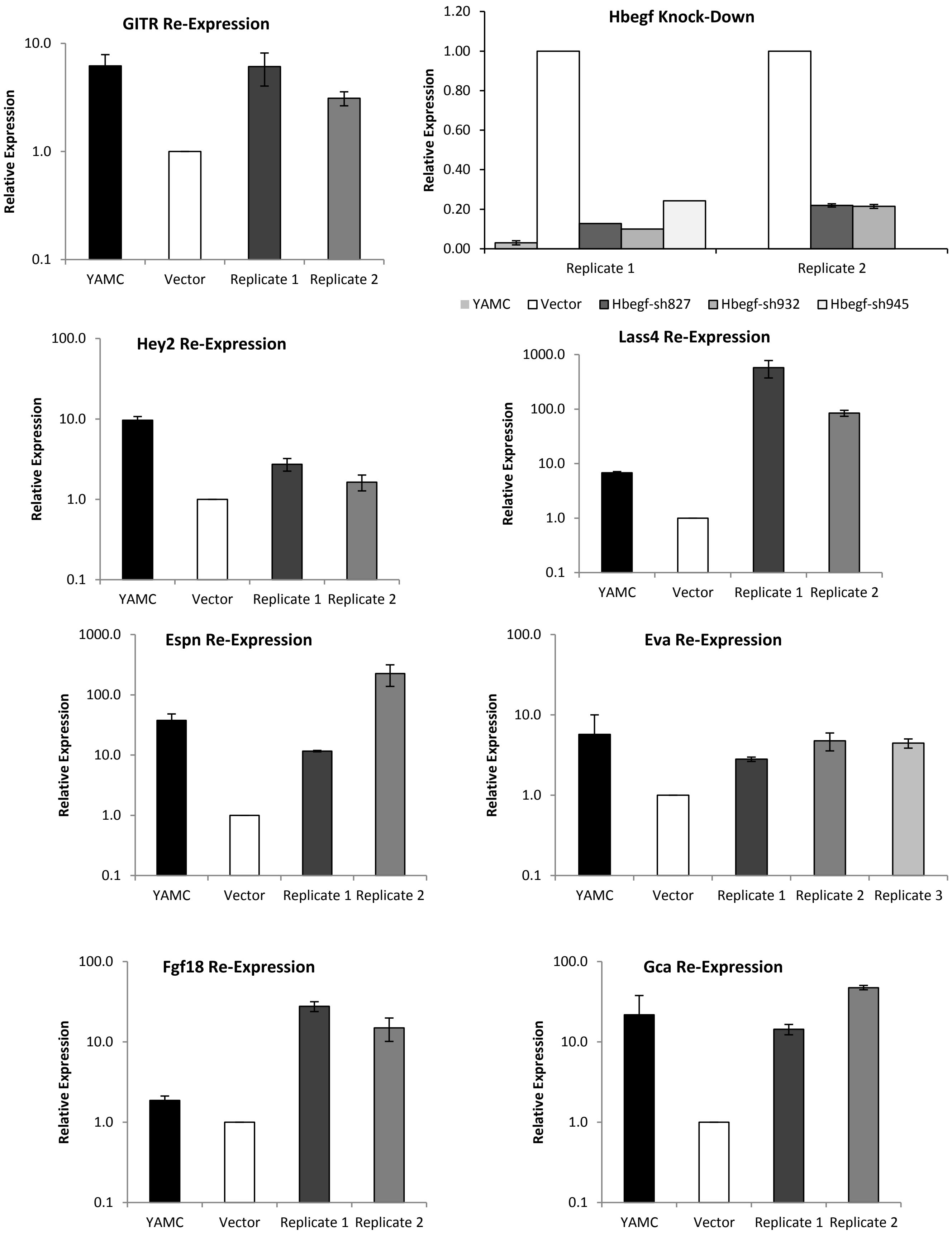


Supplemental Figure 2, continued

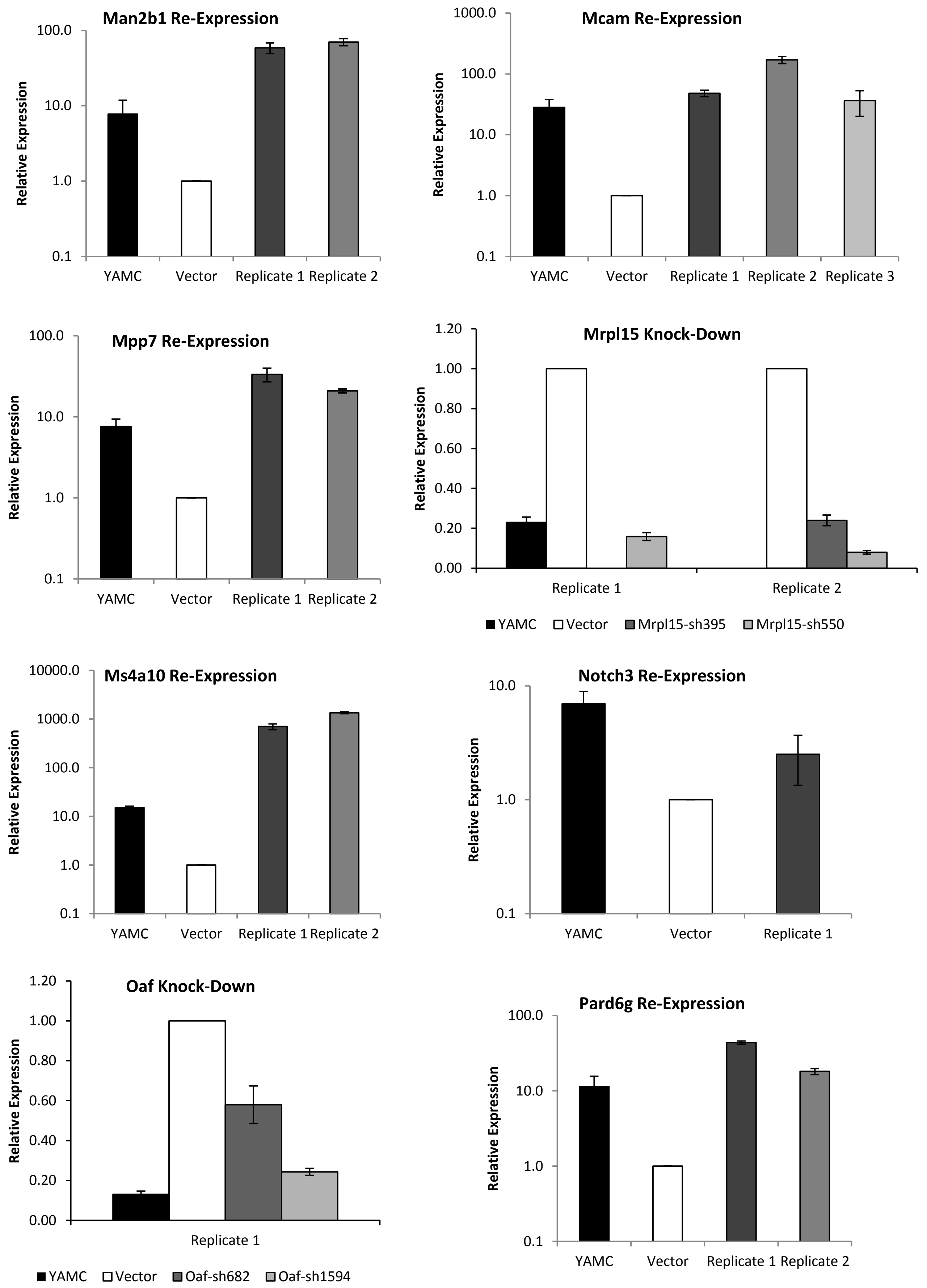


Supplemental Figure 2, continued

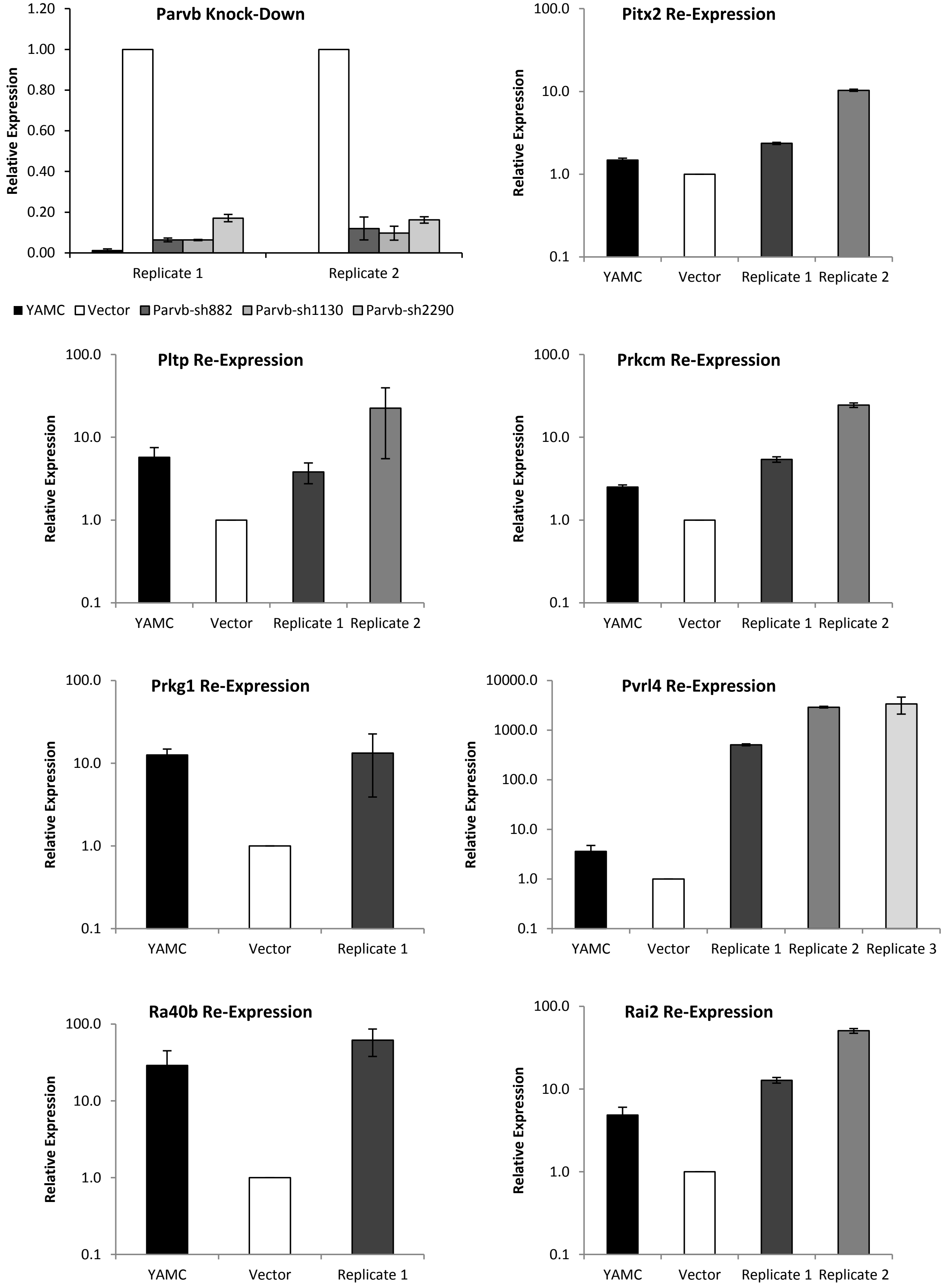


Supplemental Figure 2, continued

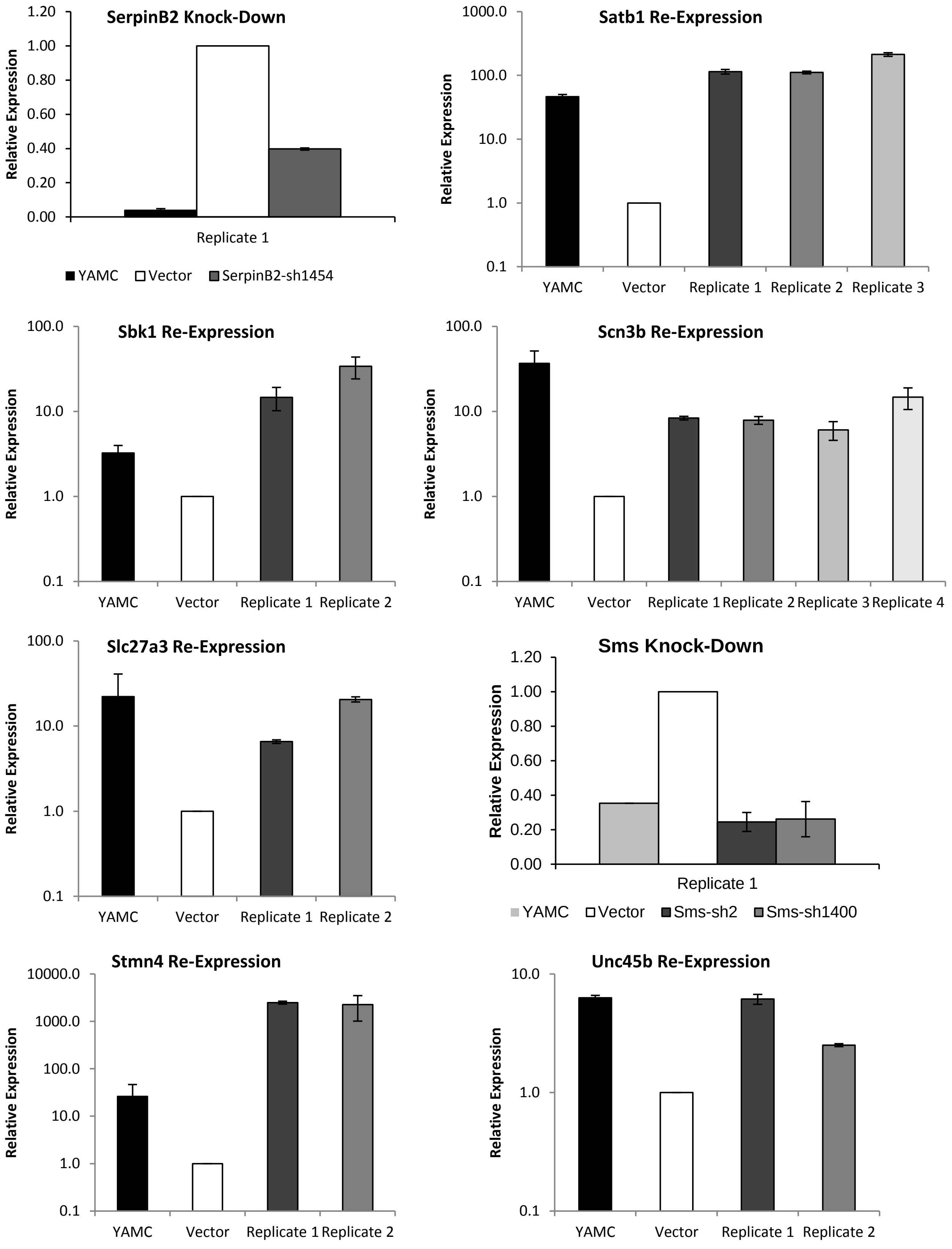


**Supplemental Figure 2: Resetting mRNA expression levels in mp53/Ras cells to levels approximating mRNA levels in non-transformed, parental YAMC cells via gene perturbations.** Each panel shows the relative expression levels of an individual gene following its perturbation in mp53/Ras cells, together with its expression levels in the matching vector control mp53/Ras cells and the parental YAMC cells, as measured by SYBR Green QPCR. Error bars indicate standard deviation of triplicate samples. Independent derivations of the perturbed cells and controls are shown individually, indicated as Replicate 1, Replicate 2, etc.

A

| Parent | Child | Direction |
|---|---|---|
| Dapk1 | Id2 | -1 |
| Dapk1 | Notch3 | 1 |
| Dapk1 | Plac8 | -1 |
| Dapk1 | Sfrp2 | 1 |
| Dapk1 | Zfp385 | 1 |
| Dffb | Hoxc13 | 1 |
| Dffb | Id2 | -1 |
| Dffb | Perp | 1 |
| Dffb | Plac8 | -1 |
| Dffb | Sfrp2 | -1 |
| Dffb | Sms | -1 |
| Dffb | Zfp385 | 1 |
| Fas | Dapk1 | -1 |
| Fas | Hoxc13 | 1 |
| Fas | Jag2 | -1 |
| Fas | Perp | 1 |
| Fas | Rgs2 | -1 |
| Fas | Sms | -1 |
| Hoxc13 | Dapk1 | -1 |
| Hoxc13 | Id2 | -1 |
| Hoxc13 | Pla2g7 | -1 |
| Hoxc13 | Plac8 | -1 |
| Hoxc13 | Rgs2 | -1 |
| Hoxc13 | Sfrp2 | 1 |
| Hoxc13 | Sms | -1 |
| Hoxc13 | Wnt9a | -1 |
| Hoxc13 | Zfp385 | -1 |
| Id2 | Rgs2 | -1 |
| Id4 | Rgs2 | -1 |
| Jag2 | Dapk1 | -1 |
| Notch3 | Dffb | 1 |
| Notch3 | Fas | 1 |
| Notch3 | Pard6g | 1 |
| Notch3 | Perp | -1 |
| Notch3 | Pla2g7 | -1 |
| Notch3 | Rab40b | 1 |
| Notch3 | Sfrp2 | 1 |
| Noxa | Dapk1 | 1 |
| Noxa | Hoxc13 | 1 |
| Noxa | Notch3 | 1 |
| Noxa | Plac8 | -1 |
| Noxa | Zfp385 | 1 |
| Pard6g | Id4 | -1 |
| Pard6g | Pla2g7 | 1 |
| Perp | Dapk1 | -1 |
| Perp | Hoxc13 | 1 |
| Perp | Id4 | 1 |
| Perp | Sms | -1 |
| Perp | Wnt9a | 1 |
| Pla2g7 | Plac8 | -1 |
| Pla2g7 | Rab40b | 1 |
| Plac8 | Wnt9a | -1 |
| Rab40b | Id4 | -1 |
| Rab40b | Notch3 | -1 |
| Rab40b | Sfrp2 | 1 |
| Rgs2 | Dapk1 | -1 |
| Rgs2 | Sfrp2 | -1 |
| Rprm | Hoxc13 | 1 |
| Rprm | Id2 | -1 |
| Rprm | Plac8 | -1 |
| Rprm | Sfrp2 | -1 |
| Rprm | Sms | -1 |
| Sms | Id4 | -1 |
| Wnt9a | Sfrp2 | -1 |

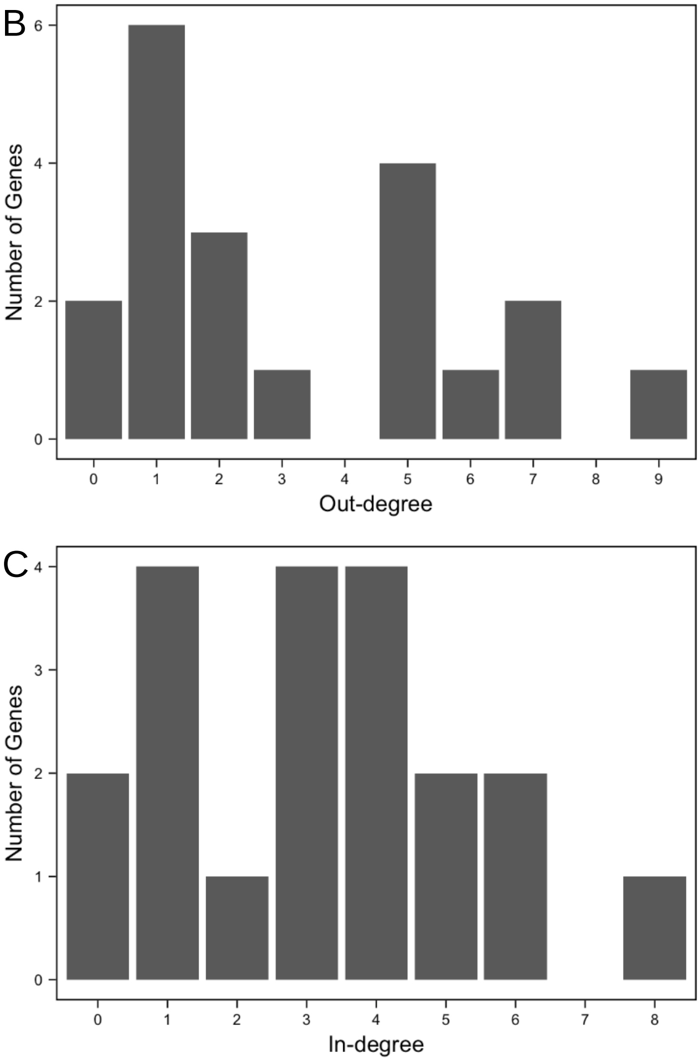


**Supplemental Figure 3: Connectedness of CRGs selected for gene regulatory network reconstruction.** A) Table indicating CRG parent-child relationships and the direction of change in the affected gene deduced from gene expression data (see supplemental file 1) following gene perturbations. B) Histogram showing the number of genes with a given out-degree, i.e. the number of children influenced by a given gene based on expression measurements. C) Histogram showing the number of genes with a given in-degree, i.e. the number of parents that influence the expression of a given gene based on expression measurements.

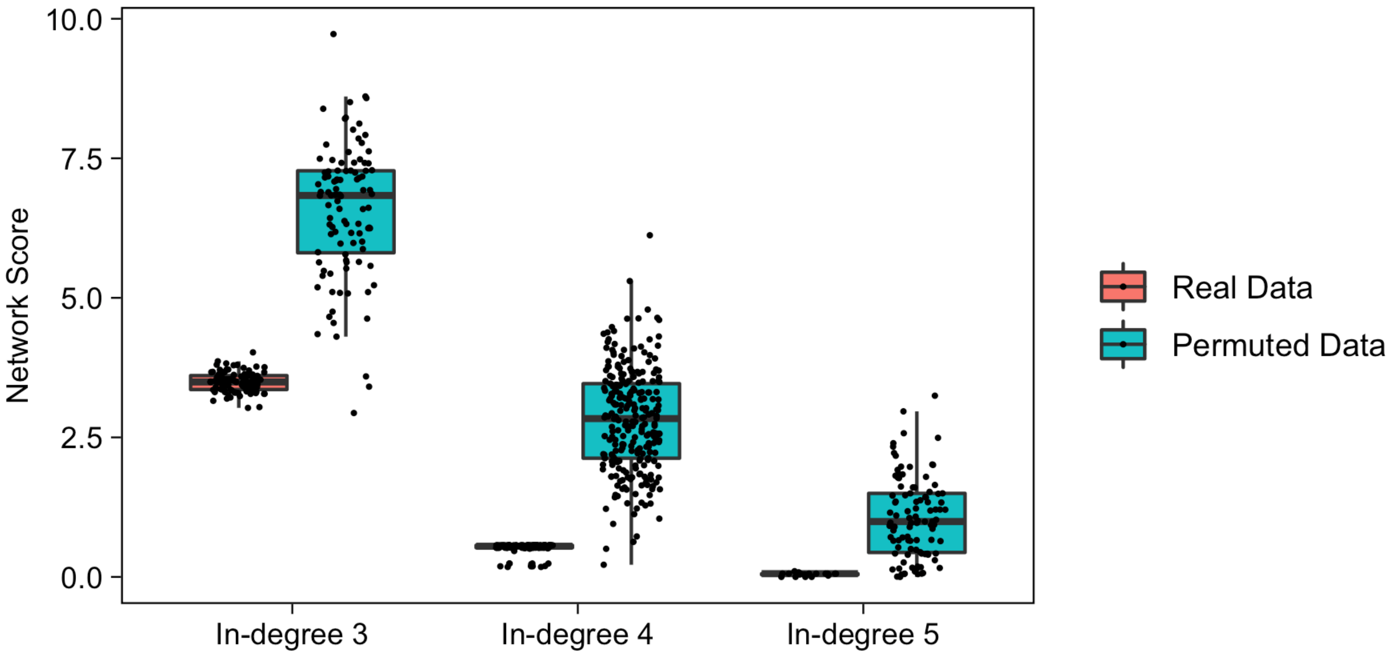


**Supplemental Figure 4: Assessment of network reproducibility and robustness.** Box plot showing network scores achieved when GRN reconstruction is performed using the true expression values (real data, salmon) as compared to permuted sets of those values (permuted data, turquoise). Comparisons for the network calculated with a maximum of three, four or five parents for each child (in-degree 3, 4 or 5) are shown.

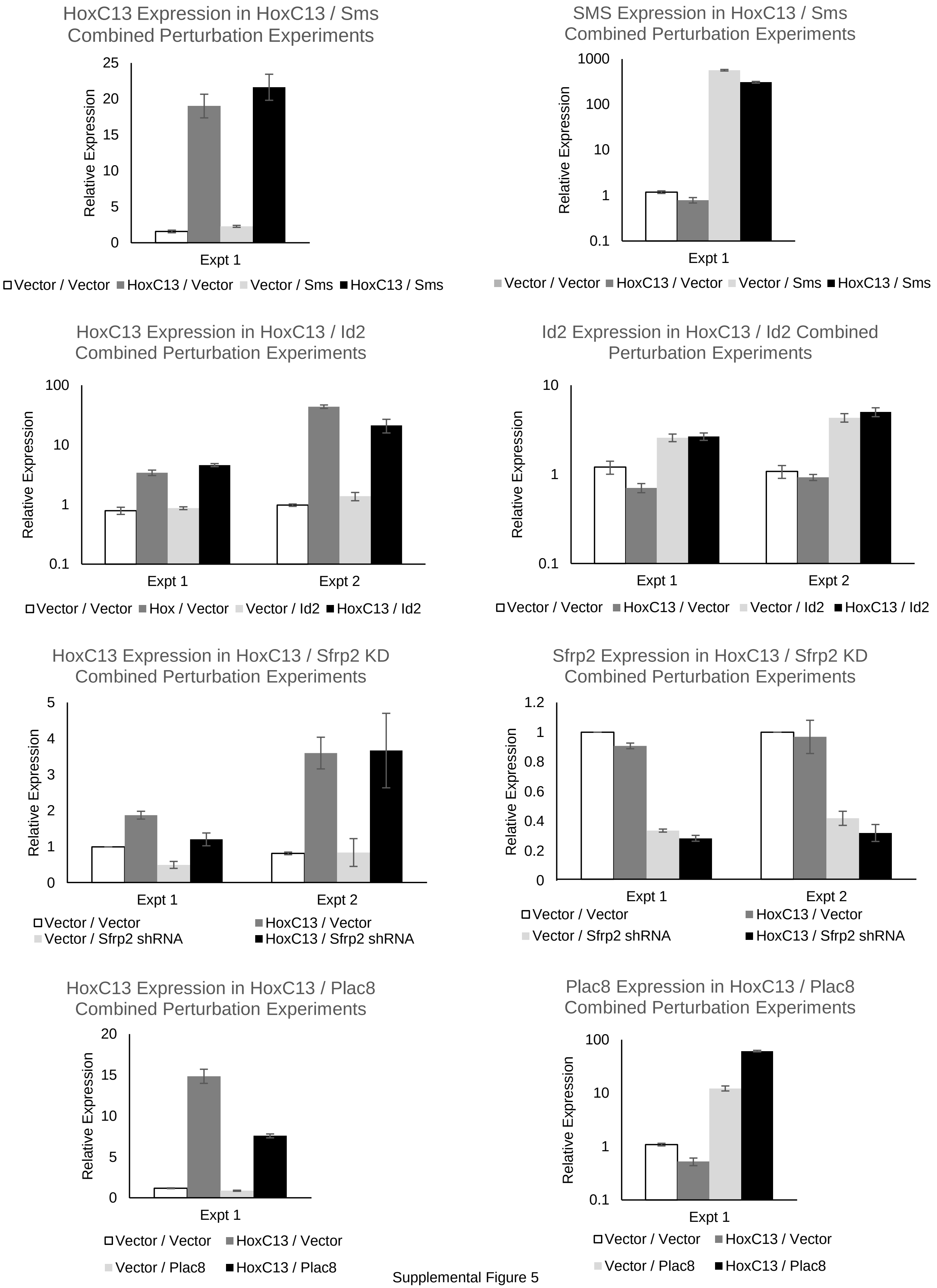


Supplemental Figure 5

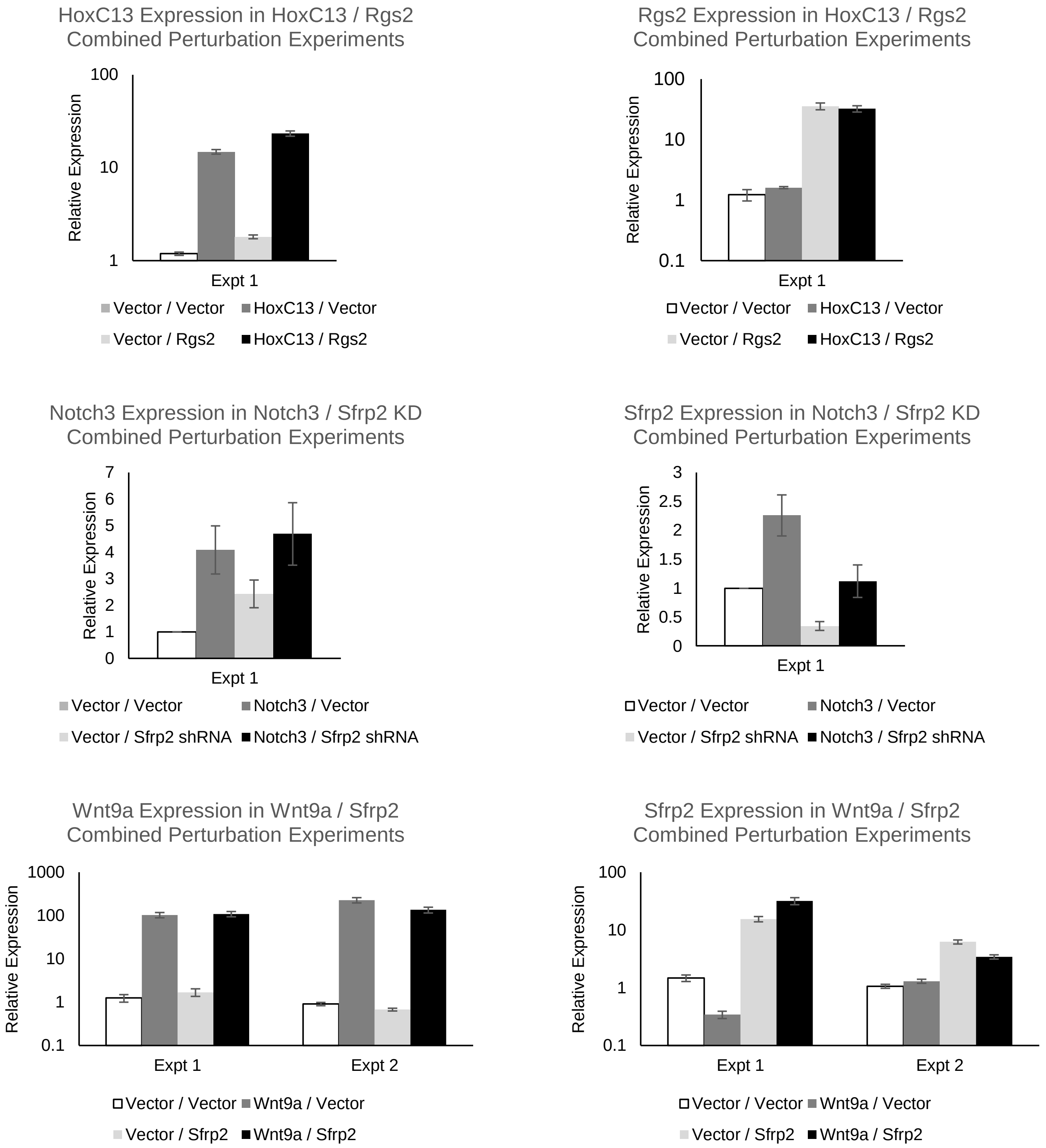


**Supplemental Figure 5: mRNA expression of the perturbagens in combinatorial gene perturbations.** Each panel shows the relative expression levels of two genes, indicated, following perturbation of one or both in mp53/Ras cells, as measured by SYBR Green QPCR. Error bars indicate standard deviation of triplicate samples. Independent derivations of the perturbed cells and controls are shown individually, indicated as Expt 1 and Expt 2, as applicable.
