## Supplemental File 1 for "Gene network modeling via TopNet reveals robust epistatic interactions between functionally diverse tumor critical mediator genes": attractor-functions.R

```
## update network state based on transition functions
updateNet <- function(state, grph, trans, pert){
  newstate <- vector(length=length(state))
  indeg <- length(dim(trans)) - 1
  for(k in 1:length(state)){
    if(k %in% pert) newstate[k] = state[k] else {
      gtmp <- grph[1:indeg, k]
      newstate[k] <- trans[matrix(c(k,state[gtmp]), ncol=indeg+1)]
    }
  }
  return(newstate)
}

findAttractorPath <- function(pgene, pstate, grph, trans, nOutcomes=3, maxIter=100){
  cond <- TRUE
  s0 <- rep(2, ncol(grph))
  s0[pgene] <- pstate
  states <- matrix(s0, ncol=1)
  while(cond){
    snew <- updateNet(states[,ncol(states)], grph, trans, pert=pgene)
    if(any(apply(states, 2, function(x) identical(x, snew)))) cond <- FALSE
    states <- cbind(states, snew)
    if(ncol(states) > maxIter) cond <- FALSE
  }
  colnames(states) <- paste0("s",0:(ncol(states)-1))
  return(states)
}

summarizeAttractor <- function(path){
  irep <- which(apply(path[,-ncol(path), drop=FALSE], 2, function(x) identical(x, path[,ncol(path)])))
  f <- function(p){
    if(all(p==2)) return(2)
    if(all(p<3)) return(1)
    if(all(p>1)) return(3)
    return(2)
  }
  apply(path[,irep:ncol(path)], 1, f)
}

scoreAttractorSummary <- function(ssObj, attrSum){
  tmp <- abs((ssObj+2) - attrSum)
  tmp[tmp>1] <- 1
  sum(tmp)
}

checkNetworkScore <- function(net, tab, verbose=TRUE){
  g0 <- t(net$parents + 1)
  t0 <- net$outcomes + 2
  netscore <- 0
  for(k in 1:(max(tab$i_exp)+1)){
    tabtmp <- tab[tab$i_exp==(k-1),]
    pgene <- tabtmp$i_node[tabtmp$is_perturbation==1]+1
    pstate <- tabtmp$outcome[tabtmp$is_perturbation==1]+2
    path <- findAttractorPath(pgene, pstate, g0, t0, maxIter=1000)
    attr <- summarizeAttractor(path)-2
    scores <- vector(length=length(attr))
    for(i in 1:length(attr)){
      scores[i] <- tabtmp$value[tabtmp$i_node==(i-1) & tabtmp$outcome==attr[i]]
    }
    netscore <- netscore + sum(scores)
  }
  if(verbose){
    cat(paste("Reported Score:", net$unnormalized_score, 
              "\nVerified Score:", netscore, 
              "\nIdentical:", identical(netscore, net$unnormalized_score)))
  } else{
    return(identical(netscore, net$unnormalized_score))
  }
}

make2Darray <- function(m){
  cbind(
    rbind(m[,,1,1], m[,,2,1], m[,,3,1]),
    rbind(m[,,1,2], m[,,2,2], m[,,3,2]),
    rbind(m[,,1,3], m[,,2,3], m[,,3,3])
  )
}

plot2Darray <- function(m2d, pnames, gname){
  require(RColorBrewer)
  tmp <- apply(m2d, c(1,2), function(x) sum(as.numeric(strsplit(x, "/")[[1]])*c(-1,0,1)))
  par(mar = c(2, 7, 7, 2))
  image(z=t(tmp[9:1,]), col=bluered(100), zlim=c(-1,1), axes=FALSE)
  abline(v=c(0.31,0.69), h=c(0.31,0.69))
  axis(side=3, at=seq(0,1,length=9), labels=rep(c("-1","0","+1"),3), tick=FALSE, line=-1)
  axis(side=3, at=seq(0.11,0.89,length=3), labels=c("-1","0","+1"), tick=FALSE, line=1, cex.axis=2)
  text(par("usr")[1]+0.05, par("usr")[4]+0.05, labels=pnames[2], xpd=TRUE)
  text(mean(par("usr")[1:2]), par("usr")[4]+0.2, labels=pnames[4], xpd=TRUE, cex=2)
  axis(side=2, at=seq(0,1,length=9), labels=rep(c("+1","0","-1"),3), tick=FALSE, las=2, line=-1)
  axis(side=2, at=seq(0.11,0.89,length=3), labels=c("+1","0","-1"), tick=FALSE, line=1, cex.axis=2, las=2)
  text(par("usr")[1]-0.06, par("usr")[4], labels=pnames[1], xpd=TRUE)
  text(par("usr")[1]-0.2, mean(par("usr")[3:4]), labels=pnames[3], xpd=TRUE, cex=2, srt=90)
  text(par("usr")[1]-0.1, par("usr")[4]+0.15, srt=45, labels=gname, xpd=TRUE, cex=3)
}
```
