## Supplemental File 1 for "Gene network modeling via TopNet reveals robust epistatic interactions between functionally diverse tumor critical mediator genes": calculate_all_probs.R

```
calculate_all_probs <- function(z){

  ## EM algorithm parameters
  tol <- 0.01 # error tolerance
  iterMax <- 500 # max iterations
  ll <- vector(length=iterMax) # log-likelihood 
  
  ## initial values
  params <- list(pneg=rep(0.25,nrow(z)),
                 p0=rep(0.5,nrow(z)),
                 ppos=rep(0.25,nrow(z))
                 )
  
  ## upper and lower limits for Uniform distributions
  Spos <- apply(z, 1, function(x) max(max(x,na.rm=T)+(max(x,na.rm=T)/sum(!is.na(x))),5))
  Sneg <- apply(z, 1, function(x) min(min(x,na.rm=T)+(min(x,na.rm=T)/sum(!is.na(x))),-5))

  ## weights for each mixture component
  wneg <- w0 <- wpos <- matrix(nrow=nrow(z), ncol=ncol(z))

  ## start EM loop
  iter <- 1
  cond <- TRUE
  while(cond){
    ## E-Step
    for(i in 1:nrow(wneg)){
      for(j in 1:ncol(wneg)){
        denom <- params$pneg[i]*dunif(z[i,j],Sneg[i],0) + params$p0[i]*dnorm(z[i,j],0,1) + params$ppos[i]*dunif(z[i,j],0,Spos[i])
        wneg[i,j] <- params$pneg[i]*dunif(z[i,j],Sneg[i],0) / denom
        w0[i,j] <- params$p0[i]*dnorm(z[i,j],0,1) / denom
        wpos[i,j] <- params$ppos[i]*dunif(z[i,j],0,Spos[i]) / denom
      }
    }
    ## M-Step
    ind <- which(!is.na(z),arr.ind=TRUE)
    params$pneg <- sapply(rowSums(wneg,na.rm=T) / rowSums(wneg+w0+wpos,na.rm=T), function(x) min(0.25,max(0.1,x)))
    params$ppos <- sapply(rowSums(wpos,na.rm=T) / rowSums(wneg+w0+wpos,na.rm=T), function(x) min(0.25,max(0.1,x)))
    params$p0 <- 1-(params$pneg+params$ppos)
  
    ## log likelihood
    ll[iter] <- sum(rowSums(wneg,na.rm=T)*(log(params$pneg)-log(-Sneg))) + 
                sum(rowSums(wpos,na.rm=T)*(log(params$ppos)-log(Spos))) + 
                sum(rowSums(w0,na.rm=T)*(log(params$p0)-0.5*log(2*pi))) - 
                sum(rowSums(w0*(z^2),na.rm=T)*(0.5))
    
    if(iter>1) cond <- (abs(ll[iter]-ll[iter-1]) > tol) & (iter < iterMax)
    iter <- iter+1
  }

 # probs <- matrix(nrow=nrow(w0),ncol=ncol(w0))
#  ipos <- which(wpos>0,arr.ind=T)
#  ineg <- which(wneg>0,arr.ind=T)
#  probs[ipos] <- wpos[ipos]
#  probs[ineg] <- -wneg[ineg]
#  rownames(probs) <- rownames(z)
#  colnames(probs) <- colnames(z)
  rownames(wneg) <- rownames(w0) <- rownames(wpos) <- rownames(z)
  colnames(wneg) <- colnames(w0) <- colnames(wpos) <- colnames(z)
  
#  probs <- probs[,order(colnames(probs))]
  return(list(pneg=wneg, p0=w0, ppos=wpos))
}
```
