## Supplemental File 1 for "Gene network modeling via TopNet reveals robust epistatic interactions between functionally diverse tumor critical mediator genes": calculate_ddCt.R

```
calculate_ddCt <- function(datNorm){
  i.ctl <- which(pData(datNorm)$sampleType=="Control")
  ddCt <- nds <- matrix(nrow=nrow(datNorm), ncol=ncol(datNorm)-length(i.ctl))
  rownames(ddCt) <- rownames(nds) <- rownames(datNorm)
  colnames(ddCt) <- colnames(nds) <- colnames(datNorm)[-i.ctl]
  
  for(k in 1:ncol(ddCt)){
    ind <- which(colnames(datNorm)==colnames(ddCt)[k])
    ind2 <- which(pData(datNorm)$batchID==pData(datNorm)$batchID[ind] & pData(datNorm)$sampleType=="Control")
    if(length(ind)!=1) stop("length(ind) does not equal 1.")
    if(length(ind2)!=1) stop("length(ind2) does not equal 1.")
    ddCt[,k] <- exprs(datNorm)[,ind] - exprs(datNorm)[,ind2]
  
    pertND <- which(featureCategory(datNorm)[,ind]=="Imputed")
    ctlND <- which(featureCategory(datNorm)[,ind2]=="Imputed")
    if(length(pertND)>0 & length(ctlND)>0) bothND <- intersect(pertND,ctlND)
  
    nds[,k] <- "none"
    if(length(pertND)>0) nds[pertND,k] <- "perturbation"
    if(length(ctlND)>0) nds[ctlND,k] <- "control"
    if(length(pertND)>0 & length(ctlND)>0) nds[bothND,k] <- "both"
  }
  
  tab.ddCt <- pData(datNorm)[pData(datNorm)$sampleName%in%colnames(ddCt),]

  SummarizedExperiment(assays=SimpleList(ddCt=ddCt, nondetects=nds),
                       colData=DataFrame(tab.ddCt))
}
```
