## Supplemental File 1 for "Gene network modeling via TopNet reveals robust epistatic interactions between functionally diverse tumor critical mediator genes": calculate_zscores.R

```
calculate_zscores <- function(ddCt){
  ## remove Rgs2 up/down perturbations and Satb1 and EphB2
  i.rm <- which(colData(ddCt)$sampleType%in%c("Rgs2:-1/1","Satb1:1","EphB2:1"))
  ddCt <- ddCt[,-i.rm]
  
  ## combine replicate perturbations that map to the same control
  SB <- as.character(paste(colData(ddCt)$batchID,colData(ddCt)$sampleType,sep=";"))
  uSB <- unique(SB)
  ddCtAvg <- matrix(nrow=nrow(ddCt),ncol=length(uSB))
  for(k in 1:ncol(ddCtAvg)){
    ind <- which(SB==uSB[k])
    ddCtAvg[,k] <- rowMeans(assay(ddCt, 1)[,ind,drop=FALSE])
  }
  rownames(ddCtAvg) <- rownames(ddCt)
  colnames(ddCtAvg) <- uSB
  
  tmp <- strsplit(uSB,";")
  tabAvg <- data.frame(sampleType=unlist(lapply(tmp, function(x) x[2])),
                       batchID=unlist(lapply(tmp, function(x) x[1])))
  
  ## compute t-statistics for each gene / perturbation
  ## shrink variances within gene
  mu <- t(apply(ddCtAvg, 1, function(x) by(x, tabAvg$sampleType, mean)))
  s2 <- t(apply(ddCtAvg, 1, function(x) by(x, tabAvg$sampleType, var)))
  n <- t(apply(ddCtAvg, 1, function(x) by(x, tabAvg$sampleType, length)))
  
  ## shrink variances
  s2shrink <- squeezeVar(s2, n-1, robust=TRUE)
  
  ## compute z-scores
  z <- mu/sqrt(s2shrink$var.post/n)
  
  ## remove the response of the perturbed gene to its own over-expression 
  ## these are sometimes nonsense due to differences between cDNA and actual transcript sequence
  for(k in 1:nrow(z)){
    ind <- grep(paste0(rownames(z)[k],":1"),colnames(z))
    z[k,ind] <- NA
  }
  
  return(z)
}
```
