## Supplemental File 1 for "Gene network modeling via TopNet reveals robust epistatic interactions between functionally diverse tumor critical mediator genes": check_matched_controls.R

```
check_matched_controls <- function(datNorm){
  i.ctl <- which(pData(datNorm)$sampleType=="Control")
  d.ctl <- d.match <- matrix(nrow=length(i.ctl), ncol=ncol(datNorm)-length(i.ctl))
  rownames(d.ctl) <- colnames(datNorm)[i.ctl]
  colnames(d.ctl) <- colnames(datNorm)[-i.ctl]
  for(i in 1:nrow(d.ctl)){
    tmp <- exprs(datNorm)[,rownames(d.ctl)[i]] 
    for(j in 1:ncol(d.ctl)){
      d.ctl[i,j] <- dist(rbind(tmp,exprs(datNorm)[,colnames(d.ctl)[j]]))
      d.match[i,j] <- pData(datNorm)$batchID[pData(datNorm)$sampleName==colnames(d.ctl)[j]] == pData(datNorm)$batchID[pData(datNorm)$sampleName==rownames(d.ctl)[i]]
    }
  }

  boxplot(d.ctl, ylab="Euclidean Distance", xaxt="n", xlab="Perturbed Samples")
  points(x=rep(1:ncol(d.ctl)), y=d.ctl[which(d.match)], pch=20, col="red")
  legend("topleft", pch=20, col="red", "matching control", cex=1.5)
}
```
