## Supplemental File 1 for "Gene network modeling via TopNet reveals robust epistatic interactions between functionally diverse tumor critical mediator genes": examine_residuals.R

```
examine_residuals <- function(ddCt, plot=FALSE){
  resids <- matrix(FALSE,nrow=nrow(ddCt), ncol=ncol(ddCt))
  for(i in 1:nrow(ddCt)){
    for(j in 1:ncol(ddCt)){
      ind <- which(colData(ddCt)$sampleType==colData(ddCt)$sampleType[j])
      resids[i,j] <- assay(ddCt,1)[i,j]-mean(assay(ddCt,1)[i,ind])
    }
  }

  if(plot){
    boxes <- list("both\nobserved"=resids[which(assay(ddCt, 2)=="none",arr.ind=TRUE)], 
                  "perturbation\nnon-detect"=resids[which(assay(ddCt, 2)=="perturbation",arr.ind=TRUE)], 
                  "control\nnon-detect"=resids[which(assay(ddCt, 2)=="control",arr.ind=TRUE)], 
                  "both\nnon-detects"=resids[which(assay(ddCt, 2)=="both",arr.ind=TRUE)])
    boxplot(boxes)
  }
  
  return(resids)
}
```
