## Supplemental File 1 for "Gene network modeling via TopNet reveals robust epistatic interactions between functionally diverse tumor critical mediator genes": format_network_input.R

```
format_network_input <- function(probs){
  
  ## organize pObj and ssObj objects for ternarynet input
  pObj <- matrix(rep(0, length(probs)), ncol=ncol(probs))
  rownames(pObj) <- rownames(probs)
  colnames(pObj) <- colnames(probs)
  ssObj <- probs
  
  for(k in 1:ncol(probs)){
    tmp <- unlist(strsplit(unlist(strsplit(colnames(ssObj)[k],"/")),":"))
    ind <- grep(tmp[1],rownames(ssObj))
    pObj[ind,k] <- ssObj[ind,k] <- as.numeric(tmp[2])
    if(length(tmp)==4){
      ind2 <- grep(tmp[3],rownames(ssObj))
      pObj[ind2,k] <- ssObj[ind2,k] <- as.numeric(tmp[4])
    }
  }
  
  ssObj <- apply(ssObj,c(1,2),as.numeric)
  pObj <- apply(pObj,c(1,2),as.numeric)
  
  return(list(pObj=pObj,ssObj=ssObj))
}
```
