## Supplemental File 1 for "Gene network modeling via TopNet reveals robust epistatic interactions between functionally diverse tumor critical mediator genes": format_network_input_scores.R

```
format_network_input_scores <- function(networkInputData){
  ngene <- nrow(networkInputData$pObj)
  nexp <- ncol(networkInputData$pObj)
  
  ## set up score data frame
  scores <- data.frame(
    i_exp=rep(0:(nexp-1), each=ngene*3),
    i_node=rep(rep(0:(ngene-1),nexp), each=3),
    outcome=rep(c(-1,0,1), nexp*ngene),
    value=rep(NA, nexp*ngene*3),
    is_perturbation=rep(0, nexp*ngene*3)
  )

  ## add which genes are perturbed
  ipert <- which(networkInputData$pObj != 0, arr.ind=TRUE)
  for(k in 1:nrow(ipert)){
    ind <- which((scores$i_exp==(ipert[k,2]-1) & scores$i_node==(ipert[k,1]-1)) 
                 & scores$outcome==networkInputData$pObj[ipert][k])
    scores$is_perturbation[ind] <- 1
  }

  ## add scores for each outcome
  for(i in 1:ngene){
    for(j in 1:nexp){
      ind1 <- which((scores$i_exp==(j-1) & scores$i_node==(i-1)) & scores$outcome==-1)
      scores$value[ind1] <- 1 + networkInputData$ssObj[i,j]
      ind2 <- which((scores$i_exp==(j-1) & scores$i_node==(i-1)) & scores$outcome==0)
      scores$value[ind2] <- abs(networkInputData$ssObj[i,j])
      ind3 <- which((scores$i_exp==(j-1) & scores$i_node==(i-1)) & scores$outcome==1)
      scores$value[ind3] <- 1 - networkInputData$ssObj[i,j]
      m <- min(scores$value[c(ind1,ind2,ind3)])
      scores$value[c(ind1,ind2,ind3)] <- scores$value[c(ind1,ind2,ind3)] - m
    }
  }

  scores$is_perturbation <- as.integer(scores$is_perturbation)
  scores$outcome <- as.integer(scores$outcome)
  
  return(scores)
}
```
