## Supplemental File 1 for "Gene network modeling via TopNet reveals robust epistatic interactions between functionally diverse tumor critical mediator genes": model_prep.R

```
model_prep <- function(dat, plot=FALSE){
    ctl <- dat[,which(is.na(colData(dat)$pGene1))]

    i.hk <- which(rownames(dat)=="Becn1")
    ctl <- ctl[-i.hk,]

    ## probability of a non-detect as a function of avg gene expression
    ## looking only at controls
    pND <- apply(assay(ctl,1),1,function(x) mean(x>39.99))
    gavg <- apply(assay(ctl,1),1,median)
    fit <- glm(pND~gavg,family=binomial(link=logit),weights=rep(ncol(ctl),length(pND)))

    if(plot){
        plot(x=gavg, y=pND, xlab="Average Expression", ylab="Proportion of non-detects",
             main="Genes across Controls", pch=20)
        xs <- seq(20, 45, by=0.01)
        lines(x=xs, y=predict(object=fit, newdata=data.frame(gavg=xs), type="response"), lwd=2)
    }
    
    ## array effects -- estimated by Becn1
    ## constrained to sum to zero
    becn1 <- assay(dat,1)[i.hk,]
    dj <- becn1 - mean(becn1)

    ## make qPCRset object
    ft <- rep("Target",nrow(dat))
    ft[rownames(dat)=="Becn1"] <- "Endogenous Control"

    fc <- matrix("OK",nrow=nrow(dat),ncol=ncol(dat))
    fc[which(assay(dat,1)>39.9999,arr.ind=TRUE)] <- "Undetermined"
    colnames(fc) <- colnames(dat)
    rownames(fc) <- rownames(dat)

    fl <- matrix("Passed", nrow=nrow(dat), ncol=ncol(dat))
    fl[which(assay(dat,1)>39.9999,arr.ind=TRUE)] <- "Flagged"
    colnames(fl) <- colnames(dat)
    rownames(fl) <- rownames(dat)

    e <- assay(dat,1)
    colnames(e) <- colData(dat)$sampleName
    
    object <- new("qPCRset", exprs=e, flag=fl)
    featureNames(object) <- rownames(dat)
    featureType(object) <- ft
    featureCategory(object) <- as.data.frame(fc)

    perts <- paste(colData(dat)$pGene1, ":", colData(dat)$dGene1, "/",
                   colData(dat)$pGene2, ":", colData(dat)$dGene2, sep="")
    perts <- gsub("/NA:NA", "", perts, fixed=T)
    perts <- gsub("NA:NA", "Control", perts, fixed=T)

    tab <- data.frame(sampleName=colData(dat)$sampleName,
                      sampleType=perts,
                      batchID=colData(dat)$batchID)
    phenoData(object) <- AnnotatedDataFrame(data=tab)

    return(list("object"=object, "fit"=fit, "dj"=dj))
}
```
