## Supplemental File 1 for "Gene network modeling via TopNet reveals robust epistatic interactions between functionally diverse tumor critical mediator genes": topology.R

```
topology <- function(fits){
  ngene <- nrow(fits[[1]]$parents)
  nexp <- length(fits[[1]]$trajectories)
  grph <- matrix(0, nrow=ngene, ncol=nexp)
  for(k in 1:length(fits)){
    gtmp <- t(fits[[k]]$parents + 1)
    for(j in 1:ncol(gtmp)){
      grph[j, gtmp[,j]] <- grph[j, gtmp[,j]] + 1
    }
  }
  cgrph = grph / length(fits)
  return(cgrph)
}
```
