## Supplemental File 1 for "Gene network modeling via TopNet reveals robust epistatic interactions between functionally diverse tumor critical mediator genes": crgnet.html

###### Matthew N. McCall

#### 2020-10-06

### Introduction

In this vignette we reproduce the analyses and results described in *Gene network modeling via TopNet reveals robust epistatic interactions between functionally diverse tumor critical mediator genes.* This package begins with a SummarizedExperiment object containing the raw qPCR Ct values from a series of perturbation experiments and corresponding control experiments. Each step in the analysis is briefly described and carried out by one or more functions implemented in this package. Each of these functions has its own help file describing it in detail; here we focus on the analysis as a whole. To avoid any confusion due to naming conventions (qPCR-based expression estimates have been called Ct values, Crt values, and Cq values to name a few), we refer to the reported Ct values as *expression estimates* or simply *expression*.

### Experimental Design

The current data consists of 66 genes (including one house keeping gene Becn1) measured across 152 samples -- 40 controls and 112 perturbations.

### Data Import and Exploration

We first load the package and data.

```
library('crgnet')
```

```
data("crgdata")
```

The Ct values from each of the samples and corresponding annotation are stored in a SummarizedExperiment object:

```
show(crgdata)
```

```
## class: SummarizedExperiment 
## dim: 66 152 
## metadata(0):
## assays(1): ct
## rownames(66): Abat Abca1 ... Wnt9a Zfp385
## rowData names(0):
## colnames(152): 1 2 ... 151 152
## colData names(6): sampleName batchID ... dGene1 dGene2
```

```
str(assay(crgdata, "ct"))
```

```
##  num [1:66, 1:152] 31.9 25.6 25.9 27 34.1 ...
##  - attr(*, "dimnames")=List of 2
##   ..$ : chr [1:66] "Abat" "Abca1" "Ank" "Ankrd1" ...
##   ..$ : chr [1:152] "1" "2" "3" "4" ...
```

```
colData(crgdata)
```

```
## DataFrame with 152 rows and 6 columns
##              sampleName     batchID      pGene1      pGene2      dGene1
##             <character> <character> <character> <character> <character>
## 1           AP.Sms.1400          AP         Sms          NA          -1
## 2     AP.Sms.1400.11.08    AP 11-08         Sms          NA          -1
## 3            AP.Sms.Sh2          AP         Sms          NA          -1
## 4       AP.Sms.Sh2.Set2     AP Set2         Sms          NA          -1
## 5             AP.Vector          AP          NA          NA          NA
## ...                 ...         ...         ...         ...         ...
## 1481 LN1.Rgs2Exp.Rgs2KD         LN1        Rgs2          NA        -1/1
## 1491         LN2.Vector         LN2          NA          NA          NA
## 150          LN2.Rgs2KD         LN2        Rgs2          NA          -1
## 151         LN2.Rgs2Exp         LN2        Rgs2          NA           1
## 152  LN2.Rgs2Exp.Rgs2KD         LN2        Rgs2          NA        -1/1
##         dGene2
##      <integer>
## 1           NA
## 2           NA
## 3           NA
## 4           NA
## 5           NA
## ...        ...
## 1481        NA
## 1491        NA
## 150         NA
## 151         NA
## 152         NA
```

The colData contains sample annotation, including the batch in which the experiment was performed, which gene(s) were perturbed, and the direction of perturbation.

#### Technical variability between samples and Becn1 as a control gene

We begin by examining the distribution of expression in each of the 40 controls.

```
controlSamples <- assay(crgdata,1)[,which(is.na(colData(crgdata)$pGene1))]
boxplot(controlSamples, xaxt="n", xlab="Control Samples")
becn1 <- controlSamples[rownames(controlSamples)=="Becn1",]
points(becn1, pch=20, col="red")
legend("bottom", pch=20, col="red", "Becn1")
```

Note that there is substantial variability in the distribution of expression across control samples. While some variability might be ascribed to the different control vectors, it is unlikely that the effect would be of this magnitude and consistency. Moreover, note that while there is substantial variability in the location of the expression distribution across control samples, the range (e.g. IQR) remains relatively constant (with a few exceptions). The house-keeping gene, Becn1, appears to track the lower quantile of the distribution of expression across the controls reasonably well. This suggests that normalizing to this house-keeping gene may perform well in these data.

```
normFactor <- becn1 - mean(becn1)
controlSamplesNormalized <- controlSamples - rep(normFactor, each=nrow(controlSamples))
boxplot(controlSamplesNormalized, xaxt="n", xlab="Control Samples Normalized")
```

The distribution of expression in the Becn1 normalized control samples, does appear to be more consistent across samples. It is unclear whether the remaining variability is due to the use of different vector controls, technical variability, or biological variability.

We can further examine the suitability of Becn1 for normalization by looking at the median absolute deviation (MAD) vs median expression.

```
controlMedians <- apply(controlSamples, 1, median)
controlMADs <- apply(controlSamples, 1, mad)
plot(x=controlMedians, y=controlMADs, pch=20, ylab="MAD", xlab="Median")
ind <- which(rownames(controlSamples)=="Becn1")
points(x=controlMedians[ind], y=controlMADs[ind], pch=20, col="red")
text(x=controlMedians[ind], y=controlMADs[ind], "Becn1", pos=4)
```

Becn1 has relatively low variability across the control samples compared to the other genes. Overall, it appears that Becn1 normalization does seem to account for much of the technical variability across samples. We will use Becn1 to normalize the data in the following section.

### Non-detect imputation and normalization

As previously reported in other data sets, we observed a strong dependency between the proportion of non-detects (those reactions failing to produce fluorescence values above a certain threshold) and the average observed expression value. Non-detects were treated as non-random missing data and imputed using the R/Bioconductor package nondetects (McCall et al. 2014).

#### Model preparation

We begin by examining the relationship between the proportion of non-detects and average expression in these data.

```
crgprepL1 <- model_prep(crgdata, plot=TRUE)
```

Additionally, the model\_prep function converts the data to the qPCRset object format needed to impute the non-detects and returns normalization factors. Specifically, the *sampleType* field of the sample annotation contains a unique description of each experiment: which gene(s) were perturbed and the direction of perturbation. This is used to define replicates for use in the imputation.

#### Non-detect imputation

We use the qpcrImpute function from the nondetects package to do the imputation. This replaces missing values (non-detects) with an imputed values based on an estimated missing data mechanism as well as the expression values seen in replicate experiments. This function takes a while to run, so the resulting data object is stored in this package and the following code is not actually evaluated in this vignette.

```
library(nondetects)
crgdataImputed <- qpcrImpute(crgprepL1$object, groupVars="sampleType")
```

To load the presaved results, we run the following:

```
data(crgdataImputed)
```

#### Normalization

We now use a function from the HTqPCR package to normalize the data to the house-keeping gene, Becn1. After normalization, we have lost the cycle thrshold (Ct) interpretation but retained the inverse relationship between expression values and the amount of transcript in the sample. Therefore, we consider the negative \(\Delta\)Ct value as our measure of normalized expression.

```
library(HTqPCR)
crgdataNorm <- normalizeCtData(crgdataImputed, deltaCt.genes="Becn1")
exprs(crgdataNorm) <- -exprs(crgdataNorm)
```

### Response to perturbation

In the previous sections we have dealt with non-detects (missing values) and normalized the data. We now turn out attention to assessing which genes are up- or down-regulated in response to each perturbation.

#### Check matched controls

First, we examine whether a given perturbed sample is more similar to the control sample from the same batch then to control samples from other batches. If there is no difference between batches, then we can compare each perturbed sample to all of the control samples. However, if there is a batch-effect, it is advantageous to compare each perturbed sample to the control sample from the same batch.

```
check_matched_controls(crgdataNorm)
```

In almost all cases, the control sample from the same batch is the most similar to the perturbed sample. This suggests that there is a difference between batches and motivates the calculation of \(\Delta\Delta\)Ct values as our measure of normalized change in expression in response to each perturbation.

#### Calculate \(\Delta\Delta\)Ct values

We calculate \(\Delta\Delta\)Ct values by computing the difference in expression between each perturbed sample and its corresponding control sample from the same batch.

```
ddCt <- calculate_ddCt(crgdataNorm)
```

#### Examine residuals

One final check of the non-detect imputation procedure from before is to examine the distribution of residuals stratified by the presence of imputed non-detect values. These can exist in either the perturbed sample, the control sample, or both samples.

```
resids <- examine_residuals(ddCt, plot=TRUE)
```

Here, we see what one would expect: a median of zero and roughly equal spread when there are no non-detects, a median slightly below zero when there is a non-detect in only the perturbed sample, and a median slightly above zero when there is a non-detect in only the control sample.

#### Probability of up- or down-regulation

In order to incorporate uncertainty into subsequent analyses, we estimate the probailitity that a gene is up- / down-regulated in response to each perturbation. We begin by calculating approximate z-scores for each perturbation:

```
zscores <- calculate_zscores(ddCt[-which(rownames(ddCt)=="Becn1"),])
```

Next we calculate the probability of up- / down-regulated in response to each perturbation by fitting a uniform / normal / uniform mixture model. This approach is similar to the probability of expression (POE) algorithm.

```
probabilities <- calculate_probs(zscores)
```

#### Filter unperturbed genes

We now filter those genes that were measured but are not perturbed in any experiment. While these genes were useful in modeling the missing data mechanism for non-detect imputation and estimating probailities of up- / down-regulation, they will not be used in the subsequent network modeling and can be removed at this point.

```
probabilities <- probabilities[-which(!rownames(probabilities)%in%gsub(":.+","",colnames(probabilities))), ]
```

Finally, we remove the six perturbation experiments that are not a single perturbation back to normal expression levels and format the probabilities for input to the network model fitting algorithm:

```
networkInputData <- format_network_input(probabilities[,-c(1,9,12,13,20,23)])
```

Both the z-scores and network input data objects are stored in this package as data objects.

#### Connectivity graph

Either the z-scores or probabilities could be thresholded to produce a connectivity graph. Here we show how to create a connectivity graph based on the probabilities used as inputs to the network modeling. We begin by showing the matrix of probabilities. Positive values denote up-regulation; negative values denote down-regulation.

```
probs <- networkInputData$ssObj
colnames(probs) <- rownames(probs)
kable(round(probs,2))
```

|  | Dapk1 | Dffb | Fas | Hoxc13 | Id2 | Id4 | Jag2 | Notch3 | Noxa | Pard6g | Perp | Pla2g7 | Plac8 | Rab40b | Rgs2 | Rprm | Sfrp2 | Sms | Wnt9a | Zfp385 |
| --- | --- | --- | --- | --- | --- | --- | --- | --- | --- | --- | --- | --- | --- | --- | --- | --- | --- | --- | --- | --- |
| Dapk1 | 1.00 | -0.11 | -0.86 | -0.52 | -0.10 | -0.15 | -0.51 | -0.19 | 0.85 | -0.10 | -0.72 | 0.18 | -0.12 | 0.08 | -0.85 | 0.07 | -0.31 | -0.11 | -0.11 | -0.11 |
| Dffb | 0.17 | 1.00 | 0.35 | -0.12 | 0.16 | -0.07 | 0.13 | 0.53 | 0.17 | 0.21 | 0.25 | -0.07 | -0.07 | 0.18 | -0.10 | 0.16 | 0.19 | 0.13 | -0.07 | 0.14 |
| Fas | 0.06 | 0.08 | 1.00 | -0.24 | 0.07 | -0.06 | 0.06 | 0.97 | 0.08 | -0.07 | 0.07 | 0.06 | -0.06 | 0.06 | -0.15 | 0.06 | 0.06 | 0.07 | 0.06 | -0.06 |
| Hoxc13 | 0.45 | 1.00 | 0.96 | 1.00 | 0.19 | -0.08 | -0.09 | 0.17 | 0.96 | -0.08 | 0.76 | 0.16 | -0.10 | -0.08 | -0.07 | 0.81 | 0.20 | 0.19 | 0.16 | 0.16 |
| Id2 | -0.78 | -0.52 | -0.19 | -0.97 | 1.00 | -0.22 | -0.17 | 0.08 | -0.48 | 0.08 | -0.18 | -0.17 | -0.28 | -0.17 | -0.20 | -0.50 | 0.08 | -0.16 | -0.17 | -0.27 |
| Id4 | -0.04 | 0.15 | 0.09 | 0.08 | 0.08 | 1.00 | 0.11 | 0.24 | -0.05 | -1.00 | 0.91 | -0.06 | 0.08 | -1.00 | -0.05 | 0.19 | -0.03 | -0.99 | 0.10 | -0.03 |
| Jag2 | -0.06 | -0.10 | -0.74 | -0.12 | 0.10 | 0.06 | 1.00 | -0.07 | 0.12 | 0.16 | -0.07 | 0.06 | -0.06 | 0.12 | -0.10 | -0.06 | 0.07 | 0.09 | 0.07 | 0.08 |
| Notch3 | 0.98 | 0.18 | -0.11 | -0.17 | 0.29 | 0.14 | 0.46 | 1.00 | 1.00 | -0.14 | 0.23 | 0.27 | 0.22 | -0.51 | -0.31 | 0.15 | 0.12 | 0.21 | 0.23 | 0.15 |
| Noxa | 0.11 | -0.11 | -0.06 | -0.07 | 0.06 | -0.08 | -0.23 | 0.11 | 1.00 | 0.06 | -0.06 | 0.06 | -0.07 | 0.10 | 0.06 | -0.10 | -0.07 | 0.09 | 0.07 | 0.42 |
| Pard6g | 0.21 | 0.08 | -0.06 | -0.14 | 0.06 | -0.07 | -0.06 | 0.71 | 0.38 | 1.00 | -0.06 | -0.06 | -0.06 | -0.08 | -0.06 | 0.06 | 0.10 | 0.16 | 0.06 | -0.06 |
| Perp | 0.31 | 0.98 | 0.99 | -0.11 | 0.11 | -0.08 | -0.15 | -0.69 | 0.10 | 0.25 | 1.00 | 0.17 | -0.13 | -0.09 | -0.08 | 0.18 | -0.06 | -0.08 | 0.11 | 0.16 |
| Pla2g7 | 0.12 | 0.08 | 0.09 | -0.81 | -0.10 | -0.13 | -0.14 | -0.81 | 0.07 | 0.51 | 0.10 | -1.00 | -0.10 | 0.08 | -0.10 | 0.25 | -0.10 | -0.13 | -0.31 | -0.26 |
| Plac8 | -1.00 | -0.57 | -0.26 | -1.00 | -0.16 | -0.16 | -0.17 | -0.17 | -0.61 | 0.21 | -0.27 | -0.72 | -1.00 | 0.12 | -0.19 | -0.96 | -0.25 | 0.11 | -0.17 | -0.17 |
| Rab40b | 0.06 | 0.07 | 0.06 | 0.06 | 0.28 | 0.07 | 0.12 | 0.64 | 0.08 | -0.30 | 0.06 | 0.58 | -0.20 | 1.00 | -0.06 | -0.07 | -0.07 | 0.06 | 0.10 | 0.06 |
| Rgs2 | 0.08 | -0.17 | -0.76 | -0.99 | -0.52 | -0.53 | -0.20 | 0.08 | 0.09 | 0.08 | 0.13 | -0.47 | -0.22 | -0.20 | -1.00 | -0.17 | 0.15 | -0.46 | -0.16 | -0.43 |
| Rprm | -0.09 | 0.06 | 0.06 | -0.11 | 0.06 | -0.07 | -0.39 | 0.09 | -0.12 | -0.15 | -0.10 | -0.06 | -0.17 | 0.08 | -0.12 | 1.00 | -0.06 | 0.06 | -0.07 | -0.06 |
| Sfrp2 | 0.98 | -1.00 | 0.18 | 1.00 | -0.26 | -0.24 | 0.25 | 0.99 | 0.18 | -0.37 | -0.14 | -0.19 | 0.32 | 0.80 | -0.80 | -1.00 | 1.00 | -0.40 | -0.85 | -0.42 |
| Sms | -0.21 | -0.99 | -0.95 | -1.00 | -0.29 | 0.07 | 0.08 | -0.17 | -0.25 | 0.09 | -0.59 | -0.20 | -0.16 | -0.23 | 0.09 | -0.73 | 0.09 | -1.00 | -0.26 | -0.19 |
| Wnt9a | 0.07 | 0.13 | 0.07 | -0.79 | 0.08 | 0.08 | -0.12 | -0.16 | -0.13 | 0.07 | 0.98 | -0.27 | -0.77 | -0.12 | 0.15 | 0.07 | -0.18 | -0.20 | 1.00 | 0.09 |
| Zfp385 | 0.82 | 0.97 | 0.23 | -0.58 | -0.13 | -0.08 | -0.08 | 0.16 | 1.00 | -0.07 | 0.38 | 0.15 | -0.25 | -0.11 | 0.15 | 0.31 | 0.15 | 0.17 | -0.08 | 1.00 |

To create a connectivity graph, we define a connection by thresholding the probabilities at an absolute value of 0.5.

```
probs[which(abs(probs)<0.5, arr.ind=TRUE)] <- 0
diag(probs) <- 0
```

Next we use the *network* package to create and plot the connectivity graph. We also add data on tumor inhibition and direction of response

```
library(network)
cgraph <- network(t(probs), matrix.type="adjacency", ignore.eval=FALSE, names.eval="probs")
data("tumor_inhibition")
set.vertex.attribute(cgraph, "tumor_inhibition", tumor_inhibition$TumorEffect)
set.edge.attribute(cgraph, "direction", sign(get.edge.attribute(cgraph,"probs")))
plot(cgraph, displaylabels=TRUE, mode="circle", boxed.labels=TRUE,
     label.bg=ifelse(get.vertex.attribute(cgraph, "tumor_inhibition")=="Smaller", "yellow", "grey"),
     edge.col=ifelse(get.edge.attribute(cgraph, "direction")==1, "red", "blue"))
legend("topleft", c("Tumor Inhibitory", "Not Tumor Inhibitory"), title="Node Color",
       fill=c("yellow","grey"))
legend("topright", c("Up-regulation", "Down-regulation"), title="Edge Color",
       col=c("red","blue"), lty=1, lwd=5)
```

### Network modeling

To assess the interdependence between these 20 CRGs, we use a ternary network model that accounts for the dynamic nature of gene regulatory networks and facilitates the evaluation uncertainty. The current methodology builds upon the modeling framework proposed in Almudevar et al. SAGMB 2011. Specifically, we assume that changes in gene expression can be modeled by one of three states – down-regulation (-1), baseline expression (0), or up-regulation (+1) – and that the state of the genes in the network at time *t+1* is a deterministic transition function of the state of the genes in the network at time *t*. This implies that from any initial state, the network will eventually reach a set of states that will repeat infinitely often, called an attractor.

Given a set of transition functions that define a network, one can easily compute the attractor for a given perturbation and assess the compatibility of that attractor with the observed steady state data. There are often many network structures that fit the observed data equally well, and the model space is huge (for the network model presented in this manuscript there are approximately 2.48×10^763 potential networks. Therefore, we use a parallel tempering algorithm (Swendsen and Wang 1986) to search the model space for networks that produce attractors that are most similar to the observed steady state data.

Unlike previous approaches, here we have incorporated uncertainty in the differential expression estimates via probabilities of up- / down-regulation. This allows the network model to give more weight to data points with higher certainty. Additionally, the ability to produce non-integer network scores eased transitions between network models and significantly decreased computational time.

Results presented in this manuscript are based on 100 independent network fits each run for 1,000,000,000 cycles in parallel across 20 processors with temperatures ranging from 0.001 to 1. The code used to produce these network fits is shown below:

```
library("pfit")
ind.arr <- as.integer(commandArgs(trailingOnly=TRUE)[1])
load("scores.rda")
results <- parallelFit(experiment_set=scores,
                       max_parents=4,
                       n_cycles=1000000000,
                       n_write=10,
                       T_lo=0.001,
                       T_hi=1,
                       target_score=0,
                       n_proc=20,
                       logfile=paste0("tnet-fit-",ind.arr,".log"),
                       seed=as.integer(ind.arr+112358)
)
```

The computational resources required to generate these network models makes in infeasible for the code to be run in this vignette. In fact the results presented in this manuscript required several weeks to generate running on the University of Rochester high performance computing cluster. However, the fits are all stored in this package for ease of use, and the source code used to generate these objects will be available in the next version of the **ternarynet** R/Bioconductor package.

### Network summaries and vizualization

Summary statistics can be computed by calculating the proportion of networks in which a given feature or features are present. One can also examine the transition functions, attractors, and trajectories all stored in the *fits* object.

#### Topology

One of the most common ways to visualize a network model is to present the topology. Here we calculate proportion of networks in which a given gene is a parent of another given gene.

```
data(networkInputData)
data(networkFits)
topo <- topology(fits)
rownames(topo) <- colnames(topo) <- rownames(networkInputData$ssObj)
kable(topo)
```

|  | Dapk1 | Dffb | Fas | Hoxc13 | Id2 | Id4 | Jag2 | Notch3 | Noxa | Pard6g | Perp | Pla2g7 | Plac8 | Rab40b | Rgs2 | Rprm | Sfrp2 | Sms | Wnt9a | Zfp385 |
| --- | --- | --- | --- | --- | --- | --- | --- | --- | --- | --- | --- | --- | --- | --- | --- | --- | --- | --- | --- | --- |
| Dapk1 | 0.00 | 0.18 | 0.01 | 0.13 | 0.37 | 0.08 | 0.09 | 0.01 | 0.98 | 0.03 | 0.72 | 0.01 | 0.20 | 0.01 | 1.00 | 0.04 | 0.01 | 0.03 | 0.06 | 0.04 |
| Dffb | 0.05 | 0.00 | 0.33 | 0.14 | 0.07 | 0.03 | 0.16 | 0.34 | 0.26 | 0.64 | 0.19 | 0.17 | 0.17 | 0.57 | 0.20 | 0.36 | 0.05 | 0.09 | 0.07 | 0.11 |
| Fas | 0.06 | 0.34 | 0.00 | 0.08 | 0.02 | 0.02 | 0.42 | 0.37 | 0.25 | 0.66 | 0.27 | 0.14 | 0.08 | 0.51 | 0.23 | 0.33 | 0.02 | 0.14 | 0.02 | 0.04 |
| Hoxc13 | 0.07 | 0.16 | 0.13 | 0.00 | 0.01 | 0.00 | 0.07 | 0.03 | 0.93 | 0.22 | 1.00 | 0.05 | 0.04 | 0.19 | 0.04 | 1.00 | 0.00 | 0.01 | 0.01 | 0.04 |
| Id2 | 0.68 | 0.33 | 0.11 | 0.83 | 0.00 | 0.00 | 0.05 | 0.15 | 0.27 | 0.09 | 0.37 | 0.06 | 0.38 | 0.08 | 0.10 | 0.14 | 0.12 | 0.08 | 0.03 | 0.13 |
| Id4 | 0.00 | 0.00 | 0.00 | 1.00 | 0.00 | 0.00 | 0.00 | 0.99 | 0.00 | 0.96 | 0.00 | 0.04 | 0.00 | 0.01 | 0.00 | 0.00 | 0.00 | 1.00 | 0.00 | 0.00 |
| Jag2 | 0.06 | 0.32 | 0.95 | 0.16 | 0.05 | 0.03 | 0.00 | 0.14 | 0.49 | 0.31 | 0.20 | 0.10 | 0.12 | 0.29 | 0.23 | 0.36 | 0.00 | 0.13 | 0.03 | 0.03 |
| Notch3 | 0.99 | 0.11 | 0.10 | 0.02 | 0.01 | 0.01 | 0.05 | 0.00 | 0.13 | 0.23 | 0.05 | 0.91 | 0.10 | 1.00 | 0.09 | 0.17 | 0.00 | 0.02 | 0.00 | 0.01 |
| Noxa | 0.10 | 0.23 | 0.34 | 0.18 | 0.10 | 0.04 | 0.32 | 0.11 | 0.00 | 0.47 | 0.14 | 0.12 | 0.21 | 0.36 | 0.41 | 0.51 | 0.01 | 0.17 | 0.09 | 0.09 |
| Pard6g | 0.19 | 0.26 | 0.34 | 0.10 | 0.07 | 0.00 | 0.16 | 0.65 | 0.36 | 0.00 | 0.17 | 0.32 | 0.17 | 0.36 | 0.17 | 0.49 | 0.02 | 0.08 | 0.01 | 0.08 |
| Perp | 0.08 | 1.00 | 0.87 | 0.05 | 0.03 | 0.01 | 0.26 | 0.28 | 0.21 | 0.34 | 0.00 | 0.08 | 0.09 | 0.29 | 0.08 | 0.18 | 0.03 | 0.03 | 0.02 | 0.07 |
| Pla2g7 | 0.11 | 0.07 | 0.16 | 0.17 | 0.09 | 0.02 | 0.05 | 0.59 | 0.13 | 0.91 | 0.21 | 0.00 | 0.08 | 0.12 | 0.24 | 0.19 | 0.08 | 0.07 | 0.08 | 0.63 |
| Plac8 | 0.55 | 0.43 | 0.08 | 0.21 | 0.52 | 0.08 | 0.02 | 0.12 | 0.47 | 0.13 | 0.05 | 0.45 | 0.00 | 0.16 | 0.04 | 0.57 | 0.01 | 0.05 | 0.02 | 0.04 |
| Rab40b | 0.08 | 0.17 | 0.20 | 0.19 | 0.09 | 0.04 | 0.12 | 0.32 | 0.22 | 0.31 | 0.04 | 0.99 | 0.15 | 0.00 | 0.42 | 0.26 | 0.02 | 0.15 | 0.08 | 0.15 |
| Rgs2 | 0.09 | 0.08 | 0.65 | 0.15 | 0.39 | 0.07 | 0.44 | 0.01 | 0.13 | 0.23 | 0.08 | 0.27 | 0.08 | 0.24 | 0.00 | 0.26 | 0.01 | 0.09 | 0.03 | 0.70 |
| Rprm | 0.09 | 0.36 | 0.35 | 0.13 | 0.11 | 0.05 | 0.24 | 0.07 | 0.62 | 0.49 | 0.13 | 0.08 | 0.27 | 0.43 | 0.38 | 0.00 | 0.02 | 0.10 | 0.03 | 0.05 |
| Sfrp2 | 0.50 | 0.00 | 0.00 | 0.99 | 0.00 | 0.00 | 0.00 | 0.99 | 0.00 | 0.00 | 0.00 | 0.00 | 0.00 | 0.01 | 0.50 | 0.00 | 0.00 | 0.01 | 1.00 | 0.00 |
| Sms | 0.25 | 0.03 | 0.23 | 0.95 | 0.27 | 0.65 | 0.11 | 0.05 | 0.49 | 0.00 | 0.05 | 0.01 | 0.07 | 0.08 | 0.14 | 0.08 | 0.04 | 0.00 | 0.46 | 0.04 |
| Wnt9a | 0.57 | 0.00 | 0.22 | 0.65 | 0.12 | 0.03 | 0.08 | 0.01 | 0.00 | 0.00 | 0.01 | 0.10 | 1.00 | 0.07 | 0.50 | 0.00 | 0.02 | 0.36 | 0.00 | 0.26 |
| Zfp385 | 0.84 | 0.58 | 0.03 | 0.40 | 0.40 | 0.00 | 0.01 | 0.09 | 0.04 | 0.07 | 0.35 | 0.14 | 0.17 | 0.08 | 0.07 | 0.65 | 0.00 | 0.03 | 0.05 | 0.00 |

We exported this information to Cytoscape to create the graphical representation of these results shown in the manuscript.

### Signficance of a good model fit

One question of interest is whether it is significant that one can obtain a low scoring network model. We examined this by permuting the network input data. For each gene, we permuted its response to all of the experiments. This retains the number of experiments to which each gene responds. In other words, the number of parents in a connectivity graph remains constant but which other genes are parents changes. We then fit a network model to the permuted data using the same parameters as above. We repeated this process 250 times.

We can now compare the network scores from the real data and the permuted data. The permuted data network scores are shown as a histogram with the real data network scores denoted with tick marks on the x-axis.

```
data(networkFits)
data(permutedNetworkFits)
real_scores <- sapply(fits, function(x) x$unnormalized_score)
permuted_scores <- sapply(pfits, function(x) x$unnormalized_score)
hist(permuted_scores, breaks=25, main="", xlab="Network Scores")
rug(real_scores, lwd=3)
```

The network scores based on the real data are less than nearly all the scores based on the permuted data (empirical p-value \(\leq 0.01\)). With the current network constraints, we can obtain good scores for the real data but not for the permuted data. This suggests that we are not extensively overfitting these data and that the current network constraints are reasonable.

### In-degree comparisons

We now examine whether we could obtain similar fits with a simpler network model. Specifically, can we reduce the number of parents (in-degree) and still fit the observed data reasonably well. To investigate this, we reran the network modeling algorithm with the *max\_parents* reduced from 4 to 3. We also considered a more complex model by increasing the *max\_parents* parameter to 5. We ran both of these models on permuted data as well.

```
data(networkFits)
data(networkFitsIndeg3)
data(networkFitsIndeg5)
indeg4_scores <- sapply(fits, function(x) x$unnormalized_score)
indeg3_scores <- sapply(fits3, function(x) x$unnormalized_score)
indeg5_scores <- sapply(fits5, function(x) x$unnormalized_score)
data(permutedNetworkFits)
data(permutedNetworkFitsIndeg3)
data(permutedNetworkFitsIndeg5)
indeg4_permuted_scores <- sapply(pfits, function(x) x$unnormalized_score)
indeg3_permuted_scores <- sapply(pfits3, function(x) x$unnormalized_score)
indeg5_permuted_scores <- sapply(pfits5, function(x) x$unnormalized_score)
plot(x=jitter(rep(c(1:6), c(length(indeg3_scores), length(indeg3_permuted_scores),
                            length(indeg4_scores), length(indeg4_permuted_scores),
                            length(indeg5_scores), length(indeg5_permuted_scores)))),
     y=c(indeg3_scores, indeg3_permuted_scores, indeg4_scores, indeg4_permuted_scores, 
         indeg5_scores, indeg5_permuted_scores),
     ylab="Network Score", xlab="", xaxt="n"
     )
axis(1, line=1.5, at=c(1.5,3.5,5.5), labels = c("In-degree 3", "In-degree 4", "In-degree 5"), 
     tick=FALSE, cex.axis=1.25)
axis(1, at=c(1:6), labels = rep(c("Real", "Permuted"), 3))
```

Increasing the in-degree cap from 3 to 4 results in a sizeable reduction in the model score; however, increasing the in-degree cap to 5 produces only a modest improvement (the in-degree 4 network models already do quite well). Regardless of the in-degree cap, better scores were achieving using the real data (as expected). The separation between real and permuted scores is greatest for an in-degree cap of 4. This lends further support to the choice of a maximum in-degree of four for these data.

### Session Info

```
sessionInfo()
```

```
## R version 4.0.2 (2020-06-22)
## Platform: x86_64-apple-darwin17.0 (64-bit)
## Running under: macOS Catalina 10.15.7
## 
## Matrix products: default
## BLAS:   /Library/Frameworks/R.framework/Versions/4.0/Resources/lib/libRblas.dylib
## LAPACK: /Library/Frameworks/R.framework/Versions/4.0/Resources/lib/libRlapack.dylib
## 
## locale:
## [1] en_US.UTF-8/en_US.UTF-8/en_US.UTF-8/C/en_US.UTF-8/en_US.UTF-8
## 
## attached base packages:
## [1] stats4    parallel  stats     graphics  grDevices utils     datasets 
## [8] methods   base     
## 
## other attached packages:
##  [1] crgnet_1.0.0                statmod_1.4.34             
##  [3] network_1.16.0              nondetects_2.18.0          
##  [5] HTqPCR_1.42.0               limma_3.44.3               
##  [7] RColorBrewer_1.1-2          SummarizedExperiment_1.18.2
##  [9] DelayedArray_0.14.1         matrixStats_0.57.0         
## [11] GenomicRanges_1.40.0        GenomeInfoDb_1.24.2        
## [13] IRanges_2.22.2              S4Vectors_0.26.1           
## [15] Biobase_2.48.0              BiocGenerics_0.34.0        
## [17] knitr_1.30                 
## 
## loaded via a namespace (and not attached):
##  [1] splines_4.0.2          gtools_3.8.2           Formula_1.2-3         
##  [4] highr_0.8              BiocManager_1.30.10    affy_1.66.0           
##  [7] latticeExtra_0.6-29    GenomeInfoDbData_1.2.3 yaml_2.2.1            
## [10] pillar_1.4.6           backports_1.1.10       lattice_0.20-41       
## [13] glue_1.4.2             digest_0.6.25          XVector_0.28.0        
## [16] checkmate_2.0.0        minqa_1.2.4            colorspace_1.4-1      
## [19] htmltools_0.5.0        preprocessCore_1.50.0  Matrix_1.2-18         
## [22] pkgconfig_2.0.3        zlibbioc_1.34.0        purrr_0.3.4           
## [25] mvtnorm_1.1-1          scales_1.1.1           jpeg_0.1-8.1          
## [28] affyio_1.58.0          lme4_1.1-23            arm_1.11-2            
## [31] tibble_3.0.3           htmlTable_2.1.0        generics_0.0.2        
## [34] ggplot2_3.3.2          ellipsis_0.3.1         nnet_7.3-14           
## [37] survival_3.2-7         magrittr_1.5           crayon_1.3.4          
## [40] evaluate_0.14          nlme_3.1-149           MASS_7.3-53           
## [43] gplots_3.1.0           foreign_0.8-80         data.table_1.13.0     
## [46] tools_4.0.2            lifecycle_0.2.0        stringr_1.4.0         
## [49] munsell_0.5.0          cluster_2.1.0          compiler_4.0.2        
## [52] caTools_1.18.0         rlang_0.4.7            nloptr_1.2.2.2        
## [55] grid_4.0.2             RCurl_1.98-1.2         rstudioapi_0.11       
## [58] htmlwidgets_1.5.2      bitops_1.0-6           base64enc_0.1-3       
## [61] rmarkdown_2.4          boot_1.3-25            gtable_0.3.0          
## [64] abind_1.4-5            R6_2.4.1               gridExtra_2.3         
## [67] dplyr_1.0.2            Hmisc_4.4-1            KernSmooth_2.23-17    
## [70] stringi_1.5.3          Rcpp_1.0.5             vctrs_0.3.4           
## [73] rpart_4.1-15           png_0.1-7              coda_0.19-4           
## [76] tidyselect_1.1.0       xfun_0.18
```
