## Supplemental File 2 for "Gene network modeling via TopNet reveals robust epistatic interactions between functionally diverse tumor critical mediator genes"

### Fitting ternery network models by replica exchange

The `retnfit` package contains a parallel implementation of the replica exchange Monte Carlo algorithm [1] (also known as parallel tempering) for fitting ternary network models [2]. Pseudocode describing the replica exchange algorithm is given below. Here,

- $\gamma$ denotes a random deviate drawn from the uniform distribution on [0,1)
- $\nu, \nu'$ denote nodes chosen at random
- $\pi(i)$ denotes one of the parents, chosen at random, of a node $i$
- $\alpha(i)$ denotes one of the outcomes, chosen at random, for the transition function of node $i$
- $\sigma$ is a possible outcome ($-1$, $0$, or $1$) chosen at random
- the cost function (the deviation of a given network from target data) is the same as described in reference 2
- For each trial move, each initial condition is advanced according to the transition functions until either a repetition is detected, or a maximum number of states is reached, in which case a large value for the cost function is assigned so that the move will not be accepted.
- the numbers of parents and transition function outcomes changed per Monte Carlo cycle is adjusted dynamically based on the fraction of those moves that are accepted, to reach a target acceptance probability of 0.5. This adjustment is done every $N_{\text{AdjustMoveInterval}}$ cycles, which is set to 7001
- exchanges between replicas are attempted every $N_{\text{ExchangeInterval}}$ cycles, which is set to 1000
- the algorithm terminates if either a target score or the maximum number of cycles is reached.

```
N_ParentMoves := 1
N_OutcomeMoves := 1
for i := 1 to N_replicas do in parallel:
    set-network-for-replica-i-to-initial-state
    T_i := T_hi (T_hi / T_lo)^((i−1)/(N_replicas−1))
for j := 1 to N_cycles do:
    for i := 1 to N_replicas do in parallel:
        if j mod 2 = 1:
            for k := 1 to N_ParentMoves
                π(ν) := ν′
        else:
            for k := 1 to N_OutcomeMoves
                α(ν) := σ
            E_i^(0) = E_i
            E_i = 0
            for k := 1 to N_InitialStates:
                set-state-to-initial-state-k
                for l := 1 to N_MaxStates
                    advance-state-k
                    if repetition-detected:
                        E_i = E_i + difference-with-target-values
                        break
                if l = N_MaxStates:
                    E_i = large-value
                    break
        if γ < exp(−[E_i − E^(0)] / T_i):
            accept-move
        else:
            restore-original-network-for-replica-i
    if j mod N_ExchangeInterval = 0:
        for i := 1 to N_replicas − 1 do in parallel:
            E_i := cost-function-for-replica-i
            E_{i+1} := cost-function-for-replica-i+1
            if γ < exp(−[E_{i+1} − E_i] [1/T_i − 1/T_{i+1}]):
                exchange-networks-for-replicas-i-and-i+1
    if best-score-for-any-replica ≤ target-score:
        break
    if j mod N_AdjustMoveInterval = 0:
        if fraction-of-parent-moves-accepted > 0.5:
            N_ParentMoves := N_ParentMoves + 1
        else if N_ParentMoves > 1:
            N_ParentMoves := N_ParentMoves − 1
        if fraction-of-outcome-moves-accepted > 0.5:
            N_OutcomeMoves := N_OutcomeMoves + 1
        else if N_OutcomeMoves > 1:
            N_OutcomeMoves := N_OutcomeMoves − 1
```
